# Metabolically purified human stem cell-derived hepatocytes reveal distinct effects of Ebola and Lassa viruses

**DOI:** 10.1101/2025.02.17.638665

**Authors:** Joseph B. Prescott, Kevin J. Liu, Angelika Lander, Nicole Min Qian Pek, Sawan Kumar Jha, Marcel Bokelmann, Manali Begur, Pang Wei Koh, Henry Yang, Bing Lim, Kristy Red-Horse, Irving L. Weissman, Kyle M. Loh, Lay Teng Ang

## Abstract

Ebola and Lassa viruses require biosafety-level-4 (BSL4) containment, infect the liver, and cause deadly hemorrhagic fevers. The cellular effects of these viruses, and whether different families of hemorrhagic-fever viruses elicit similar effects, remain fundamental questions in BSL4 virology. Here, we introduce a new metabolic selection approach to create nearly-pure hepatocytes from human pluripotent stem cells, killing non-liver cells by withholding essential nutrients. Unexpectedly, Ebola and Lassa exerted starkly different effects on human hepatocytes. Ebola infection activated the integrated stress response (ISR) and WNT pathways in hepatocytes in vitro and killed them, whereas Lassa did not. Within non-human primates, Ebola likewise infected hepatocytes and activated ISR signaling *in vivo*. In summary, we present a single-cell transcriptional and chromatin accessibility roadmap of human hepatocyte differentiation, purification, and viral infection.

## INTRODUCTION

Risk Group 4 viruses are among the deadliest viruses known on Earth and must be studied in specialized biosafety level 4 (BSL4) laboratories, few of which exist worldwide^1^. Sparingly few approved therapies exist for the devastating diseases caused by these viruses. Accordingly, many of these pathogens have been designated as World Health Organization Priority Diseases, necessitating urgent research and development^2, 3^. Filoviruses such as Ebola virus (∼25-90% fatality rate, species *Orthoebolavirus zairense*), Sudan virus (∼54% fatality rate, species *Orthoebolavirus sudanense*), and Marburg virus (∼81% fatality rate, species *Orthomarburgvirus marburgense*) cause periodic outbreaks in Africa^4-6^. These include the 2013-2016 Ebola virus epidemic that killed over 11,000 individuals in West Africa^7-10^, the 2022 Sudan virus epidemic in Uganda^11, 12^, and the 2023 Marburg virus outbreaks in Equatorial Guinea and Tanzania^13^. Lassa virus (∼1% fatality rate, species *Lassa mammarenavirus*) is an arenavirus that infects up to 500,000 individuals every year in Africa^14-16^. One important question is what are the effects of these Risk Group 4 viruses on physiologically-relevant human cell-types. Additionally, few, if any, studies have directly compared the effects of different Risk Group 4 viral families—such as filoviruses (family *Filoviridae*) vs. arenaviruses (family *Arenaviridae*)—on human cells, in the same experimental system^14, 17^. While many Risk Group 4 viruses cause deadly hemorrhagic fevers in humans, an important and unresolved question is whether distinct families of Risk Group 4 viruses cause common or diverging effects on their shared human cells.

The liver is a primary target of many deadly Risk Group 4 viruses: Ebola, Sudan, Marburg, and Lassa viruses infect liver cells *in vivo* and can ultimately cause liver damage^18-36^. Although it has been known for more than 30 years from patient autopsies that Ebola, Sudan, Marburg, and Lassa viruses infect liver cells, the unique constraints of BSL4 experimentation^37^ have made it challenging to study the mechanistic effects of these viruses on specific human cell-types, such as hepatocytes. As surrogates, liver cancer cell lines HepG2 and HuH7 have been extensively employed in BSL4 virology to study cellular responses to viral infection and to conduct chemical and genetic screens^38-43^. However, liver cancer cell lines lack fundamental liver hallmarks, harbor chromosome abnormalities^44, 45^, and are largely incapable of producing interferons, key antiviral cytokines^46-48^. Alternatively, primary hepatocytes isolated from human beings are the current gold standard for studying liver biology *in vitro*, but they are scarce, expensive, and vary between individuals. Importantly, it is well-established that primary hepatocytes swiftly lose hepatocyte identity and functions *ex vivo*^49-53^.

The ability to generate massive numbers of human hepatocytes would provide a valuable platform to study the effects of Risk Group 4 viruses on a physiologically-relevant cell-type *in vitro*. Human pluripotent stem cells (hPSCs, including embryonic and induced pluripotent stem cells) can be grown in large numbers and can theoretically generate any cell-type within the body, including hepatocytes^54-58^. However, hPSCs can generate hundreds of different cell-types, and it remains challenging to differentiate them exclusively into hepatocytes. Despite considerable progress, hPSC differentiation yields heterogeneous cell populations containing hepatocytes commingled with non-liver cells^59-81^. These contaminating non-liver cells pose challenges for virology, regenerative medicine, and other applications. For instance, one study differentiated hPSCs into a heterogeneous population comprising ∼25% ALBUMIN+ hepatocytes and ∼75% unidentified cells to study Ebola virus infection^48^. However, the predominance of non-liver cells made it unclear whether Ebola directly infected hepatocytes or whether any observed cellular responses to Ebola were attributable to hepatocytes and/or the contaminating non-liver cells^48^.

Here we develop “metabolic selection”, a new, rapid, and simple approach to purify hPSC-derived hepatocytes by selectively depleting non-liver cells. This is rooted in the emerging concept that different cell-types have distinct metabolic requirements to survive and can thus be selectively killed by withholding specific nutrients^82-87^. We hypothesized that hepatocytes could uniquely withstand deprivation of glucose and other specific nutrients, owing to their ability to break down glycogen into glucose and to exploit alternate nutrient sources for energy (gluconeogenesis)^88-91^. Indeed, we found that withholding 3 essential nutrients (glucose, pyruvate, and glutamine) for 1-3 days destroyed non-liver cells *in vitro*, while hepatocytes survived. Metabolic selection thus produces pure populations of hPSC-derived hepatocytes without recourse to surface marker-based cell sorting or other purification schemas. Critically, metabolic selection purifies hepatocytes based on their metabolic *functionality*, instead of surface marker expression^92^.

hPSC-derived hepatocytes purified by metabolic selection offer a new human hepatocyte model with several advantages for BSL4 virology relative to extant liver cancer cell lines. For instance, hPSC-derived hepatocytes behaved similarly to adult primary human hepatocytes in their ability to trigger interferon signaling, whereas liver cancer cell lines could not. Additionally, hPSC-derived hepatocytes showed greater transcriptional similarity to primary hepatocytes than liver cancer cell lines.

Having established highly pure hPSC-derived hepatocytes as an enhanced model system, we discovered that Ebola and Lassa viruses—although both causing viral hemorrhagic fevers *in vivo*—had starkly different transcriptional and cytopathic effects on human hepatocytes. Notably, Ebola killed hepatocytes, induced WNT and integrated stress response (ISR) pathway genes, and suppressed liver function genes. In contrast, Lassa virus resulted in transient transcriptional changes in genes largely different from those induced by Ebola virus, and did not induce either WNT or ISR pathways. To our knowledge, this is the first direct comparison of how Ebola and Lassa viruses affect human cells in the same experimental system, thus representing a step forward for comparative virology. We further confirmed that Ebola virus directly infects hepatocytes *in vivo* and activates ISR signaling within a non-human primate model of Ebola virus disease. Finally, we also present a single-cell transcriptional and chromatin accessibility roadmap of hPSC differentiation into hepatocytes, and how they compare with HepG2 and HuH7 liver cancer cell lines (https://anglab.shinyapps.io/liverscrna/). Taken together, the ability to create purified populations of hPSC-derived hepatocytes by metabolic selection will enable a range of applications in BSL4 virology, stem cell research, and regenerative medicine.

## RESULTS

### Stepwise changes in gene expression, chromatin accessibility, and cellular diversity during hPSC differentiation into hepatocytes

To chart a stepwise molecular roadmap for hepatocyte development, we applied single-cell RNA-sequencing (scRNAseq^93^), bulk-population RNA-seq, and OmniATAC-seq^94^ to a modified version of our previously-developed schema wherein hPSCs are efficiently differentiated into day 1 primitive streak, day 2 definitive endoderm, day 3 posterior foregut, day 6 liver bud progenitors, day 12 early hepatocytes and day 18 hepatocytes^69, 81^ (**Figs. 1A**-**B**). At each differentiation step, we provided extracellular signals to direct hPSC differentiation into a desired cell-type while inhibiting other extracellular signals to suppress the formation of unwanted fates (**Fig. 1A**). Flow cytometry confirmed that our differentiation strategy sequentially generated 99.2% pure MIXL1^+^ primitive streak, 99.8% pure CXCR4^+^ definitive endoderm, and 82.4% pure ALBUMIN^+^ hepatocytes by days 1, 2 and 18 of differentiation, respectively (**Fig. 1C**).

**Fig. 1.**
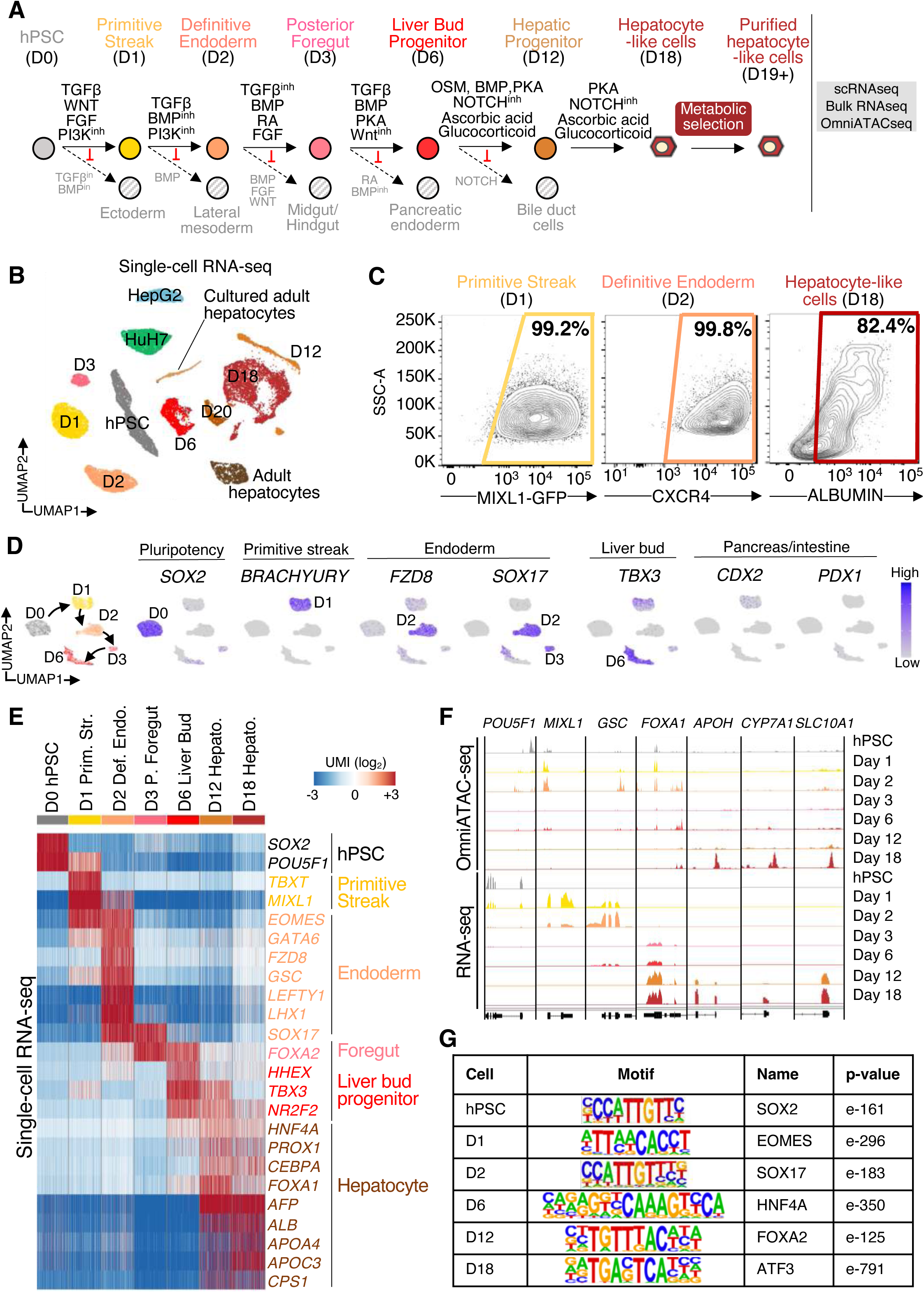
Transcriptional and chromatin roadmap for human liver differentiation from hPSCs. A) Stepwise differentiation of hPSCs towards hepatocytes modified from^69^, followed by metabolic selection to purify hepatocytes. B) scRNAseq of H1 hPSC differentiated toward hepatocytes (days 1-18 [d1-d18] of differentiation), liver cancer cell lines (HepG2 and HuH7), and freshly-thawed and cultured adult human hepatocytes. An embedding in 2D space by uniform manifold approximation and projection (UMAP), of how 12 samples and 52,678 cells was analyzed. C) Flow cytometry reveals generation of MIXL1+ anteriormost primitive streak (assayed by flow cytometry of *MIXL1-GFP* hPSCs^197^), CXCR4+ definitive endoderm, and ALBUMIN+ hepatocytes within 1, 2 and 18 days of hPSC differentiation, respectively. D) scRNAseq of H1 hPSCs differentiated toward day 6 liver bud progenitors. The entire cell population was harvested without pre-selection. Gene expression quantified in log_2_ UMI units. E) Heatmap of selected liver function genes from scRNAseq of H1 hPSCs differentiated into day 18 hepatocytes. The entire cell population was harvested without pre-selection. Each column represents the gene expression of a single cell, clustered by differentiation timepoints. F) Bulk-population RNAseq and OmniATACseq of H1 hPSCs differentiated towards hepatocytes. G) Transcription factor motifs enriched within accessible chromatin regions at each stage of H1 hPSC differentiation towards hepatocytes, as determined by OmniATACseq.

To rigorously assess the purity and synchrony of differentiation, we performed scRNAseq to systematically identify all cell-types arising at each differentiation step (**Fig. 1B, Fig. S1A**). Within 1 day of hPSC differentiation, pluripotency transcription factor (TF) *SOX2* was sharply downregulated, and primitive streak TFs (*BRACHYURY*, *MIXL1*) were uniformly expressed (**Figs. 1D, Fig. S1B-C**). Next, both day 2 definitive endoderm and day 3 posterior foregut homogeneously expressed the endodermal TF *SOX17*^95-97^, with definitive endoderm-specific markers (*FZD8*, *GSC*, *CER1*)^98-102^ expressed in day 2 endoderm but declining in day 3 foregut (**Fig. 1D, Fig. S1C**). After 6 days of differentiation, the liver bud marker *TBX3*^103^ was homogeneously expressed, without the apparent expression of intestinal or pancreatic markers (*CDX2*, *PDX1*)^104-106^, confirming high differentiation precision (**Fig. 1D, Fig. S1C**). Sub-clustering analyses suggested that day 1 primitive streak, day 2 definitive endoderm, day 3 posterior foregut, and day 6 liver bud progenitors were largely homogeneous, with residual population heterogeneity at each respective step of differentiation primarily driven by cell-cycle markers (**Fig. S1C**). This thus reveals largely uniform progression from pluripotency to primitive streak, definitive endoderm, posterior foregut, and liver bud progenitors. On days 12 and 18 of differentiation, liver bud progenitor marker *TBX3* gradually decreased, and conversely, hepatocyte markers (*ALB*, *CPS1, APOA4*, *APOC3*) were upregulated (**Fig. 1E**). These stepwise changes in gene expression were also observed by bulk-population RNA-seq (**Fig. S1D, Table S1**).

At each stage of differentiation, there were stepwise changes in chromatin accessibility, with sequential opening and closure of pluripotency, primitive streak, endodermal, and liver gene loci, often coincident with changes in gene expression (**Figs. 1E-F, S1D, Table S1**). Analysis of TF motifs enriched in cell-type-enriched accessible chromatin regions revealed sequential progression from pluripotency (POU5F1) to primitive streak (EOMES), endodermal (SOX17), and eventually liver (HNF4A) TFs, corresponding to the sequential expression of their cognate TFs (**Figs. 1G, S1B**). Taken together, this provides a rich resource to potentially discover new markers and regulators of human liver differentiation and can be accessed via an interactive browser: https://anglab.shinyapps.io/liverscrna/.

### hPSC-derived hepatocytes arise alongside a subset of intestinal cells

Having generated >80% pure ALBUMIN+ hPSC-derived hepatocytes within 18 days of differentiation, we next explored the identity of the remaining non-liver cells (**Fig. 2A**, **Fig. S2A**). We posited that non-liver cells arising during hepatocyte differentiation *in vitro* might correspond to other endodermal tissues arising near the liver *in vivo* (e.g., intestines)^107^. Indeed, ∼8% of cells co-expressed CDX2 and PDX1 (**Fig. 2A**, **Fig. S2A**), indicative of the duodenum^104-106^, which is the anterior region of the small intestine and is adjacent to the developing liver *in vivo* (**Fig. 2B**). scRNAseq identified the majority of day 18 cells as *ALBUMIN*+ hepatocytes but also discovered three smaller subsets of non-liver cells: 1) intestinal goblet-like cells (*TFF1*, *TFF3*, *CREB3L1*+^108-110^), 2) intestinal enteroendocrine-like cells (*CHGA*, *ARX*, *NEUROD1*+^111, 112^) and 3) trace numbers of mesenchymal cells (*LUM*, *COL3A1*+) (**Fig. 2C**, **Fig. S2B**). Notably, both intestinal goblet and enteroendocrine cells are specified by NOTCH blockade^113-115^, consistent with our use of NOTCH inhibitor to differentiate hPSC-derived liver bud progenitors into hepatocytes^69^. In summary, our differentiation approach generates an enriched population of hepatocytes, with some non-liver cells arising contemporaneously. These non-liver cells could interfere with studies of hepatocyte biology and viral infection, and we thus sought to eliminate them.

**Fig. 2.**
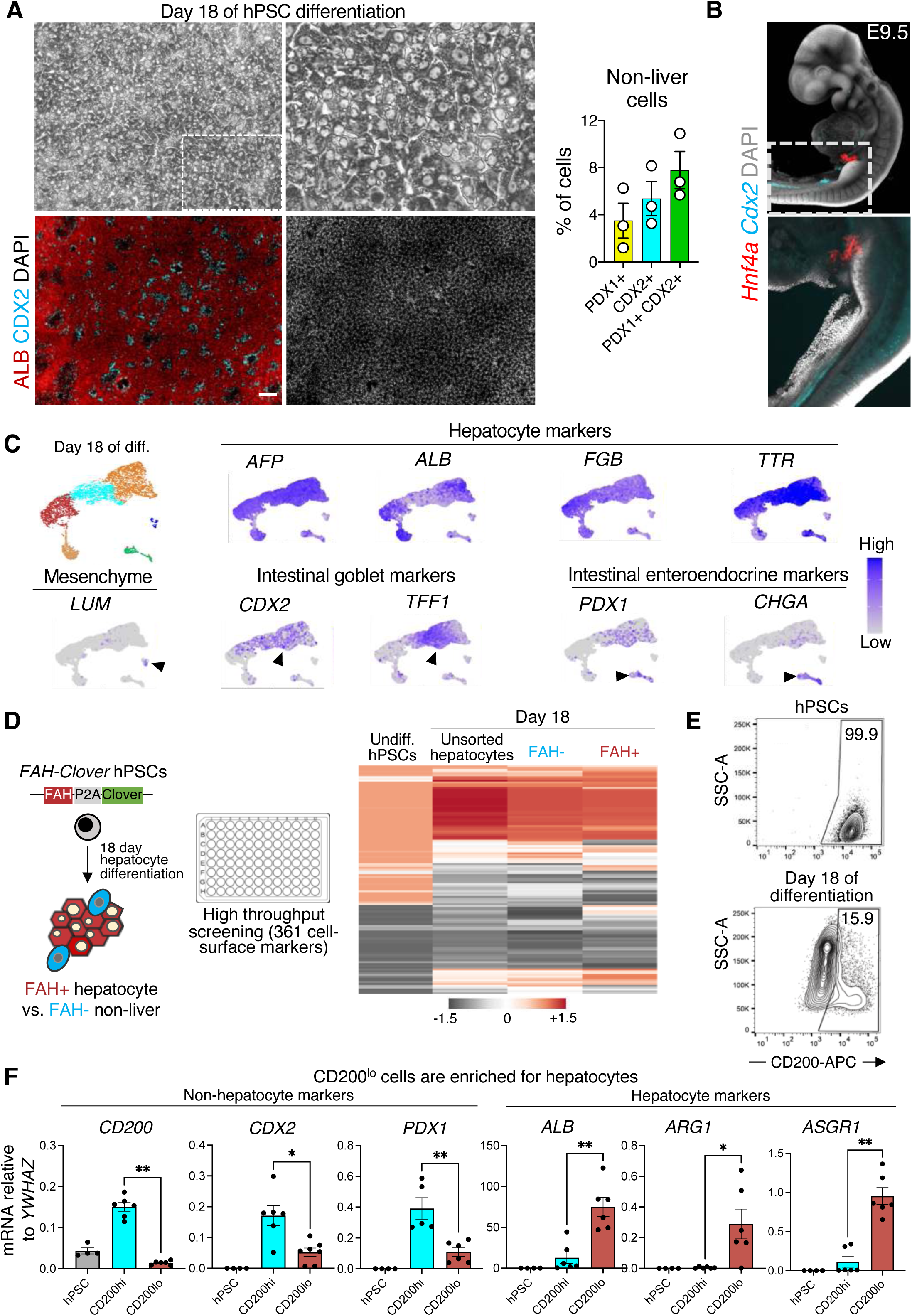
Intestinal cells emerge alongside hPSC-derived hepatocytes and can be removed using cell-surface marker CD200. A) ALBUMIN (ALB), CDX2, and PDX1 immunostaining of H1 hPSC-derived day 18 hepatocyte-containing populations, followed by image quantification. Scale bar, 100μm. Top right represents a magnification of the top left brightfield image. Error bars denote standard errors. B) *Hnf4a* and *Cdx2* mRNA staining of a mouse embryo at embryonic day 9.5 (E9.5) by HCR3.0^198^, showing the proximity of developing liver and intestines. C) scRNAseq of day 18 H1 hPSC-derived hepatocyte-containing populations, followed by Louvain clustering ^199^ to identify multiple cell-types within the population. The entire cell population was harvested without pre-selection. Gene expression quantified in log_2_ UMI units. D) High throughput antibody screening of *FAH-2A-Clover* reporter hPSCs^69^, which were either undifferentiated, differentiated for 18 days (“unsorted hepatocytes”), or day 18 populations that were subset into Clover+ vs. Clover-subpopulations. E) Flow cytometry of CD200 expression on H1 hPSCs before or after differentiation into day 18 hepatocytes. F) qPCR of CD200^hi^ vs. CD200^lo^ populations isolated from day 18 hPSC-derived hepatocyte populations by fluorescence-activated cell sorting (FACS) and undifferentiated hPSCs as a negative control. *P<0.05, **P<0.01. Error bars denote standard errors.

### Cell-surface marker CD200 distinguishes hepatocytes vs. non-liver cells

To eliminate non-liver cells, we first discovered surface markers that distinguish hepatocytes from non-liver cells. To this end, we performed a high-throughput antibody screen of 361 cell-surface markers in *FAH-2A-Clover* knock-in reporter hPSCs^69^ differentiated towards hepatocytes (**Fig. 2D, Table S2**). FAH is a tyrosine metabolic enzyme expressed in hepatocytes^116^. We found that CD200 was low in FAH+ hepatocytes but was enriched in FAH-non-liver cells *in vitro* (**Fig. 2E**). Likewise, *CD200* is expressed in 19 different human tissues but is absent from the liver and hepatocytes (**Figs. S2C-D**)^117, 118^, reaffirming that CD200 is a non-liver marker *in vivo*. *In vitro*, CD200 was highly expressed by hPSCs, definitive endoderm, and liver bud progenitors but significantly decreased by day 18, where only ∼15.9% of cells remained CD200^hi^ (**Figs. 2E, S2E**). We further tested the value of CD200 as a marker for hepatocyte purification by performing fluorescence-activated cell sorting (FACS) of CD200^hi^ vs. CD200^lo^ cells after 18 days of hPSC differentiation (**Fig. 2F, Fig. S2F**). CD200^lo^ cells were enriched for *ALB*+, *ARG1*+, and *ASGR1*+ hepatocytes, whereas CD200^hi^ cells comprised *CDX2*+ and *PDX1*+ non-liver cells (**Fig. 2F**), as confirmed by scRNAseq (**Fig. S2G**).

### Metabolic selection purifies hPSC-derived hepatocytes and destroys non-liver cells

As a parallel approach to purifying hPSC-derived hepatocytes, we developed a new “metabolic selection” strategy to selectively kill non-liver cells. Remarkably, hepatocytes are one of the few cell types capable of generating glucose *de novo*^88-91^ and might thus withstand nutrient depletion. We hypothesized that withholding glucose and other nutrients *in vitro* would eliminate most cell-types (**Fig. S3A**). Lending confidence that hepatocytes might endure nutrient starvation, we found that hPSC-derived hepatocytes expressed genes involved in energy generation from alternative sources, including genes integral to glycogen breakdown (*PGYL*, *AGL*), glycogen synthesis (*GBE1*), glutamine synthesis (*GS*), galactose metabolism (*GALK1*, *GALK2*, *GALE*, *GALT*), and gluconeogenesis (*G6PC*, *FBP1*, *PCK1*) (**Fig. 3A, Table S3**). In contrast, hPSC-derived hepatocytes expressed lower levels of glycolysis genes (*HK1/2*, *PKM1/2*) than hPSCs (**Fig. 3A**), suggesting that hepatocytes may be less dependent on glycolysis for energy production. Consistent with this, gene ontology (GO) analysis showed that hPSC-derived hepatocytes expressed a wide range of metabolic genes involved in glucose, lipid, and triglyceride metabolism (**Fig. S3B**), suggesting that they could exploit alternative substrates such as lipids and triglycerides to survive a starvation episode.

**Fig. 3.**
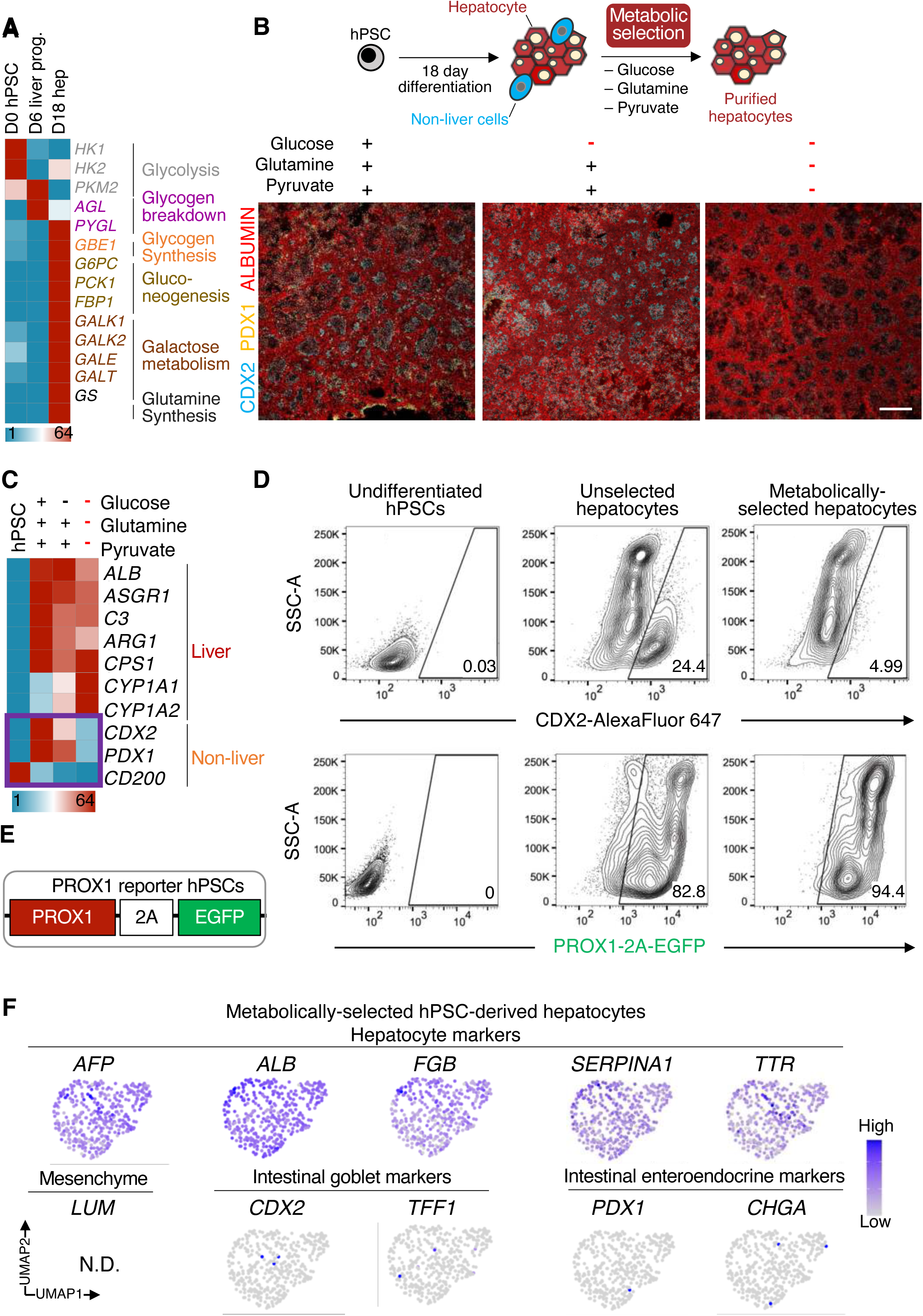
Metabolic selection eliminates non-liver cells, thereby purifying hPSC-derived hepatocytes. A) qPCR of undifferentiated H1 hPSCs, day 6 liver bud progenitors, and day 18 hepatocytes reveals the upregulation of metabolic pathway genes in hepatocytes. B) ALBUMIN, CDX2, and PDX1 immunostaining of H1 hPSC-derived day 18 hepatocytes cultured in standard vs. metabolic selection media for 3 additional days. Scale bar, 100μm. C) qPCR of H1 hPSC-derived hepatocytes cultured in standard vs. metabolic selection media for 3 days, with undifferentiated hPSCs shown as a negative control. D) Flow cytometry of undifferentiated H1 hPSCs and day 18 H1 hPSC differentiated hepatocytes stained for CDX2 in standard vs. metabolic selection media for 2 days. E) Flow cytometry of undifferentiated H1 *PROX1-2A-EGFP* hPSC reporter line and day 18 H1 *PROX1-2A-EGFP* hPSC reporter line differentiated hepatocytes in standard vs. metabolic selection media for 1 day. F) scRNAseq of H1 hPSC-derived hepatocytes cultured in metabolic selection media for 2 days reveals near-absence of non-liver marker expression. The entire cell population was harvested, without pre-selecting cell subsets. Gene expression quantified in log_2_ UMI units.

We found that withholding exogenous glucose, glutamine, and pyruvate for 1-3 days led to the enrichment of hPSC-derived hepatocytes and significantly depleted CDX2+ and PDX1+ non-liver cells (**Figs. 3B, 3D, 3F, Figs. S3C-E**). Metabolically-selected hepatocytes are >94% pure, as assayed using newly engineered PROX1-2A-EGFP hPSCs (**Fig. 3E**). This process of nutrient deprivation—which we refer to as “metabolic selection”—also destroyed undifferentiated hPSCs, day 6 liver bud progenitors, and day 6 midgut/hindgut endoderm (intestinal progenitors), while sparing day 18 hepatocytes (**Fig. S3C**). Combinatorial withdrawal of exogenous nutrients was critical: withholding glucose alone spared undifferentiated hPSCs and failed to robustly eliminate CDX2+ and PDX1+ non-liver cells (**Fig. 3B, Fig. S3D**). Consistent with how hPSCs can redundantly rely on glycolysis and glutamine oxidation for survival^119, 120^, we found that removing both glucose and glutamine destroyed a range of non-liver cells (**Fig. S3C**). However, withholding glucose, glutamine, and pyruvate most effectively eliminated non-liver cells, including hPSCs, liver bud progenitors, and midgut/hindgut cells, while sparing hPSC-derived hepatocytes (**Fig. S3C**). We term our metabolic selection media lacking all three essential nutrients (glucose, glutamine, and pyruvate) as “HepSelect” media.

scRNAseq revealed that two days of HepSelect treatment stringently purified hepatocytes. Hepatocyte markers *ALB*, *AFP*, *FGB*, *SERPINA1,* and *TTR* were homogeneously expressed after metabolic selection (**Fig. 3E**). By contrast, markers of intestinal (*CDX2*, *TFF1*, *PDX1*, *CHGA*) and mesenchymal cells (*LUM*) were essentially undetectable (**Fig. 3E**). RNA-seq and qPCR also confirmed that metabolic selection reduced the expression of non-liver markers including *CDX2*, *PDX1*, and *CD200* (**Fig. 3C**, **Fig. S3F, Table S4**). In sum, metabolic selection provides a new, effective, and scalable approach to purify hPSC-derived hepatocytes based on their metabolic functionality. Metabolic selection does not require specialized equipment; it solely entails treatment with nutrient-depleted media and is therefore potentially simpler and less expensive to apply on a large scale than other cell purification schemas.

### hPSC-derived hepatocytes are transcriptionally more similar to adult hepatocytes than liver cancer cell lines often used in BSL4 virology

scRNAseq revealed that hPSC-derived hepatocytes were more transcriptionally similar to adult hepatocytes than liver cancer cell lines HuH7 and HepG2 (**Figs. 4A-B**), which are widely used in BSL4 virology^38-43^. As controls, we used primary adult human hepatocytes (pooled from 100 individuals to reduce donor-to-donor variability) that were freshly isolated or cultured for 6 days.

**Fig. 4.**
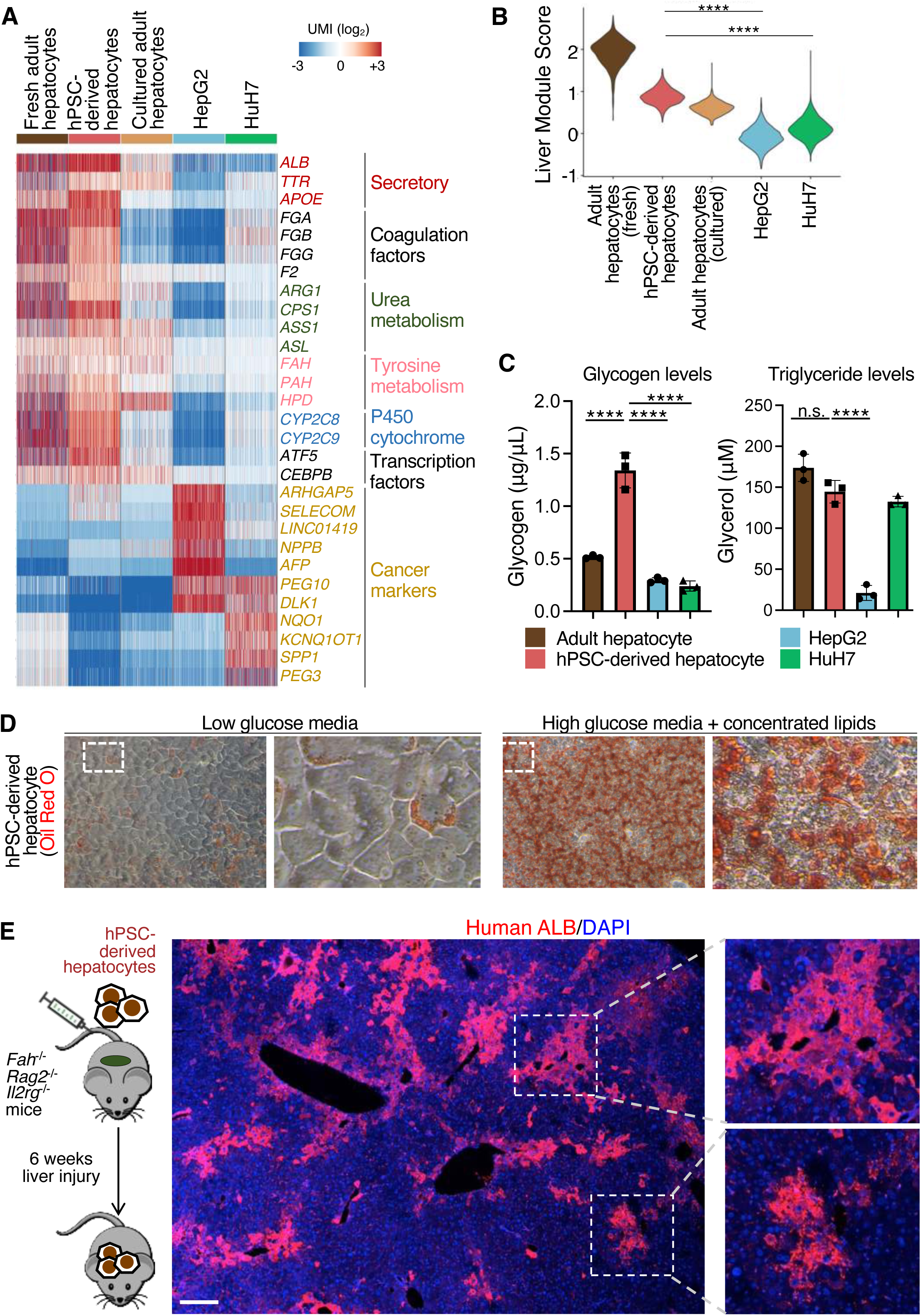
Comparison of metabolically-selected hPSC-derived hepatocytes, adult primary hepatocytes, and liver cancer cells. A) Heatmap of selected liver function genes from scRNAseq of metabolically-selected H1 hPSC-derived hepatocytes, adult primary hepatocytes that were either freshly isolated or cultured for 6 days, and liver cancer cell lines (HepG2 and HuH7). Each column represents the gene expression of a downsampled number of 300 cells, clustered together by cell-type. B) Average expression levels of prevailing hepatocyte marker signature genes^123^ across metabolically-selected H1 hPSC-derived hepatocytes, freshly-thawed and cultured adult primary hepatocytes, and liver cancer cell lines (HepG2 and HuH7), as quantified by scRNAseq. ****P<0.0001 in unpaired parametric t-test C) Quantification of glycogen and glycerol levels across metabolically-selected H1 hPSC-derived hepatocytes, cultured adult primary hepatocytes, and liver cancer cell lines (HepG2 and HuH7). Error bars denote standard deviations. D) Oil Red O staining of H1 hPSC-derived hepatocytes differentiated in either low glucose media vs. high glucose and lipid-rich media for 12 days.

Many hepatocyte genes essential for carrier protein and coagulation factor production, as well as urea, tyrosine, and xenobiotic metabolism, were highly expressed by both hPSC-derived hepatocytes and adult hepatocytes (**Fig. 4A**). However, these genes showed low or near-absent expression in HuH7 and HepG2 cells (**Fig. 4A**). Focusing on genes integral to two hallmark hepatocyte functions—detoxification of ammonia into urea (*CPS1*, *OTC*, *ASS1*, *ASL*, *ARG1*) and fibrinogen production (*FGA*, *FGB*, *FGG*)—we found that hPSC-derived hepatocytes expressed considerably higher levels of these genes than HuH7 or HepG2 cells (**Fig. 4A, Fig. S4A**). On the contrary, HuH7 and HepG2 expressed cancer markers *PEG3* and *PEG10*^121, 122^, which were otherwise absent from adult or hPSC-derived hepatocytes (**Fig. 4A**). Scoring these various liver cell models using prevailing hepatocyte marker signatures^123^ showed that metabolically-selected hPSC-derived hepatocytes scored similarly to cultured adult hepatocytes and scored higher than HuH7 and HepG2 (**Fig. 4B**). Transcriptome-wide analysis revealed that primary adult hepatocytes exhibited 79% and 81% transcriptional similarity to hPSC-derived hepatocytes and cultured adult hepatocytes, respectively (**Fig. S4B**). On the other hand, HuH7 and HepG2 cancer cells were transcriptionally less similar (67% and 69%, respectively) to adult hepatocytes (**Fig. S4B**). Taken together, hPSC-derived hepatocytes harbor transcriptional hallmarks of hepatocyte identity and function that have been lost in liver cancer cell lines.

However, hepatocytes *in vivo* and *in vitro* were not identical, and exhibited important differences. Upon culture, adult hepatocytes downregulated expression of liver markers and cytochrome enzymes *CYP2C8, CYP3A4, TAT, SERPINA3, APOC3,* and *CYP4A11*^53, 124-131^ (**Fig. S4C**), and upregulated fetal hepatocyte marker *AFP* (**Fig. S4D; Table S6**). Likewise, hPSC-derived hepatocytes *in vitro* also expressed *AFP* but lower *CYP3A4* levels than adult hepatocytes (**Fig. S4D**). In sum, hPSC-derived hepatocytes *in vitro* are more similar to primary hepatocytes relative to liver cancer cell lines, although they are not identical to primary hepatocytes *in vivo*.

### hPSC-derived hepatocytes display metabolic functions *in vitro* and engraft the mouse liver

Purified hPSC-derived hepatocytes also executed various liver metabolic functions *in vitro*, such as (1) storage of glucose in the form of glycogen and (2) conversion of free fatty acids into triglycerides^88, 89, 132-136^. hPSC-derived hepatocytes stored ∼4.5-to 5.6-fold greater glycogen than HepG2 and HuH7 cells (**Fig. 4C**). Triglyceride levels were ∼6.9-fold higher in hPSC-derived hepatocytes relative to HepG2 cells (**Fig. 4C**). Additionally, hPSC-derived hepatocytes treated with high glucose and lipids accumulated lipid droplets (**Fig. 4D**). These results also suggest glycogen and triglycerides as possible energy sources for hPSC-derived hepatocytes, allowing them to withstand nutrient deprivation in the aforementioned metabolic selection strategy.

Additionally, hPSC-derived hepatocytes could engraft the injured liver of immunodeficient *Fah*^-/-^ *Rag2*^-/-^ *Il2rg*^-/-^ (FRG) mice (**Fig. 4E**). FRG mice lack tyrosine metabolism enzyme *Fah* and thus develop chronic liver failure^137^. 6 weeks after transplantation, hPSC-derived hepatocytes repopulated the mouse liver, as shown by the abundant number of human ALBUMIN+ hepatocytes *in vivo* (**Fig. 4E**). Of note, hPSC-derived hepatocytes were spatially distributed across the liver lobule, some of which were nearby blood vessels, such as the central and portal veins (**Fig. S4E**). hPSC-derived hepatocytes adjacent to the central vein expressed the pericentral hepatocyte marker GLUTAMINE SYNTHETASE (GS)^138-140^ (**Fig. S4E**). Hence, hPSC-derived hepatocytes could engraft the injured liver, attesting to their potential for liver regeneration.

### hPSC-derived and primary hepatocytes can express interferon-stimulated genes in response to double-stranded RNA analog, unlike liver cancer cell lines

Furthermore, we explored the ability of hPSC-derived hepatocytes to activate interferon signaling, an innate immunity pathway with antiviral effects^141^. Liver cancer cell lines HuH7 and HepG2 have been widely used to study Ebola and other viruses under BSL4 containment^38-43^, yet they are defective in interferon production ^46-48^, which hampers their utility for studying viral infection *in vitro*.

One day of treatment with poly(I:C), a double-stranded RNA analog^142, 143^, strongly induced interferon-stimulated genes (ISGs) *MX1*, *VIPERIN*, *IFIT1*, *IFIT2*, *IFIT3*, *OAS1*, *OASL*, and *ISG15*^144^ in hPSC-derived hepatocytes obtained from two different hPSC lines, H1 and H7 (**Fig. 5A**). Similarly, poly(I:C) upregulated ISGs in primary adult human hepatocytes obtained from two different sources (**Fig. 5A**). As expected, poly(I:C) largely failed to upregulate these ISGs in HepG2 and HuH7 liver cancer cell lines (**Fig. 5A**)^46-48^. RNA-seq confirmed significant ISG and inflammatory cytokine upregulation in poly(I:C)-treated hPSC-derived and primary adult hepatocytes, but minimal changes in HepG2 and HuH7 cells (**Fig. 5B, Table S5**). Taken together, hPSC-derived hepatocytes behaved more similarly to primary hepatocytes in their induction of ISGs than liver cancer cell lines. Having established that hPSC-derived hepatocytes can activate interferon signaling, we next applied these cells to model infection by Risk Group 4 viruses.

**Fig. 5.**
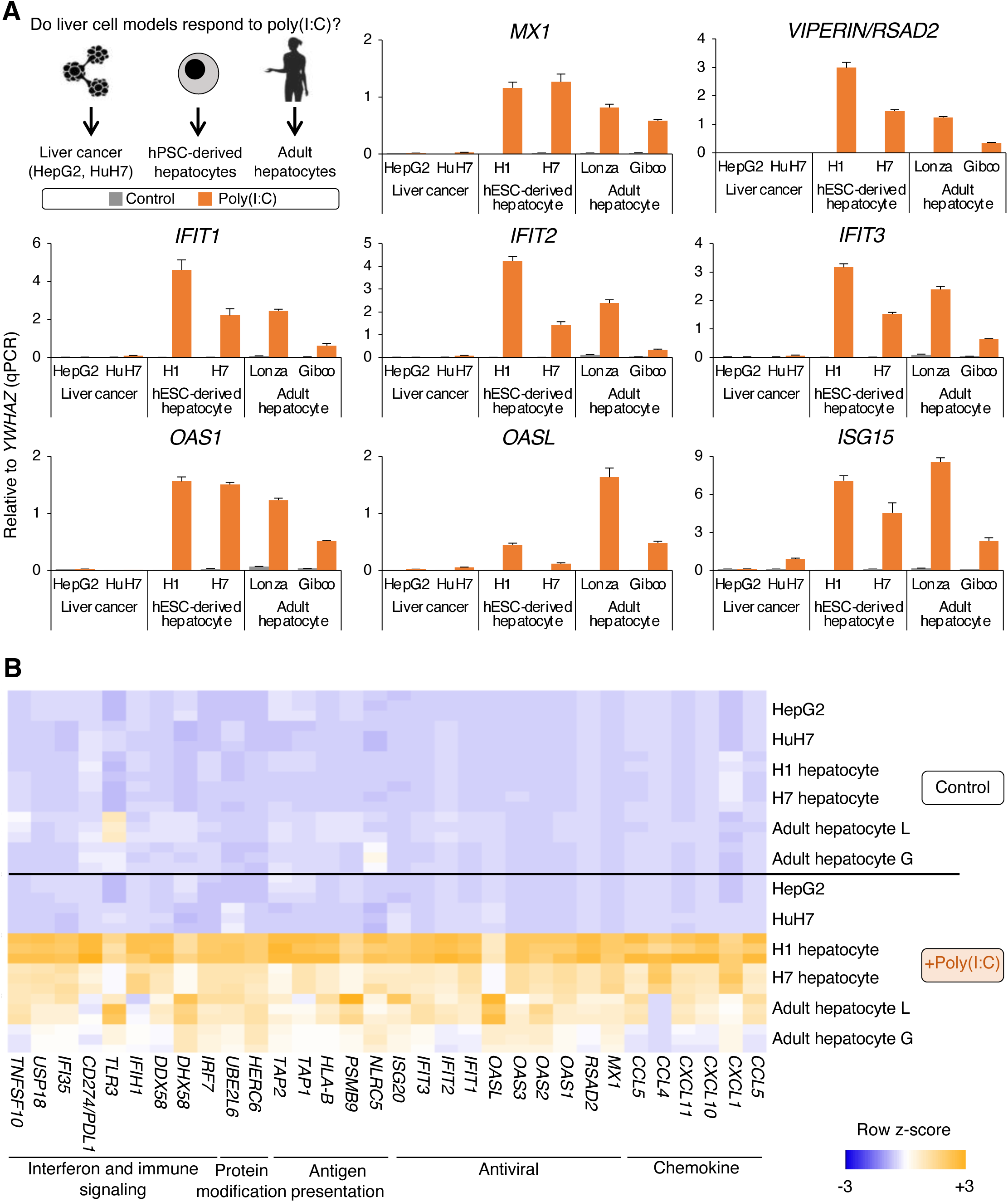
hPSC-derived and adult hepatocytes activate interferon signaling in response to poly(I:C), but liver cancer cells do not. A) qPCR of hPSC-derived hepatocytes (generated from H1 or H7 hPSC lines), adult primary human hepatocytes (procured from Lonza [L] or Gibco [G]), or liver cancer cells (HepG2 or HuH7) treated with 50 μg/mL poly(I:C) or control media for 1 day. Gene expression data normalized to *YWHAZ*, such that *YWHAZ* expression = 1.0. Expression of interferon-stimulated genes^144^ is shown. Error bars denote standard errors. B) Heatmap of selected liver function genes from RNA-seq of hPSC-derived hepatocytes (generated from H1 or H7 hPSC lines), adult primary human hepatocytes (procured from Lonza [L] or Gibco [G]) or liver cancer cells (HepG2 or HuH7) treated with 50 μg/mL poly(I:C) or control media for 1 day.

### Risk Group 4 viruses can infect purified hPSC-derived hepatocytes

The liver is an important target of many deadly Risk Group 4 viruses. It has been known for over 30 years from autopsies of infected humans and macaques that liver cells are infected by Ebola virus^21, 23, 30^, Sudan virus^21, 28, 29^, Marburg virus^19, 27^, and Lassa virus^20, 24-26^ *in vivo*.

To test whether hPSC-derived hepatocytes could be infected by Risk Group 4 viruses, we first developed methods to cryopreserve, thaw, and culture hepatocytes before viral infection under BSL4 containment. However, thawed hPSC-derived hepatocytes cultured in standard commercially-available media rapidly lost the expression of hepatocyte genes (**Fig. S5A**). We hypothesized that hepatocytes could be maintained by continued exposure to hepatocyte-specifying signals, akin to those used to specify hepatocytes from hPSCs. Indeed, continued exposure to hepatocyte-specifying signals—glucocorticoid and PKA activation and NOTCH inhibition^69^ (**Fig. S5A**), together with high-density culture (**Fig. S5B**)—maintained hPSC-derived hepatocytes. Mechanistically, PKA and high cell density activate the Hippo pathway, which is required for hepatocyte maturation^145-148^. In these conditions, thawed hPSC-derived hepatocytes could be cultured for 1 week while maintaining the expression of hallmark hepatocyte genes (**Fig. S5C**).

To visually track viral spread over time, we infected purified hPSC-derived hepatocytes with recombinant viruses wherein fluorescent proteins were inserted into authentic viral genomes: GFP-expressing Ebola virus^149^, ZsGreen-expressing Sudan virus^150^, ZsGreen-expressing Marburg virus^151^, and ZsGreen-expressing Lassa virus^43^ (**Figs. 6A**, **Figs. S6A, S6E**). Ebola, Sudan, Marburg, and Lassa virus extensively replicated in hPSC-derived hepatocytes, as shown by focus forming assays of the culture media (**Fig. 6B**) and direct fluorescent imaging of infected hepatocytes (**Fig. 6C**). We confirmed that these viruses infect hepatocytes, as cells co-expressed viral markers (ZsGreen or GFP) and the hepatocyte marker ALBUMIN (**Fig. 6D, Fig. S6C**). In sum, Ebola, Sudan, Marburg, and Lassa viruses can efficiently infect and replicate in hPSC-derived hepatocytes *in vitro*, providing a tractable platform to study the effects of these viruses on human hepatocytes.

**Fig. 6.**
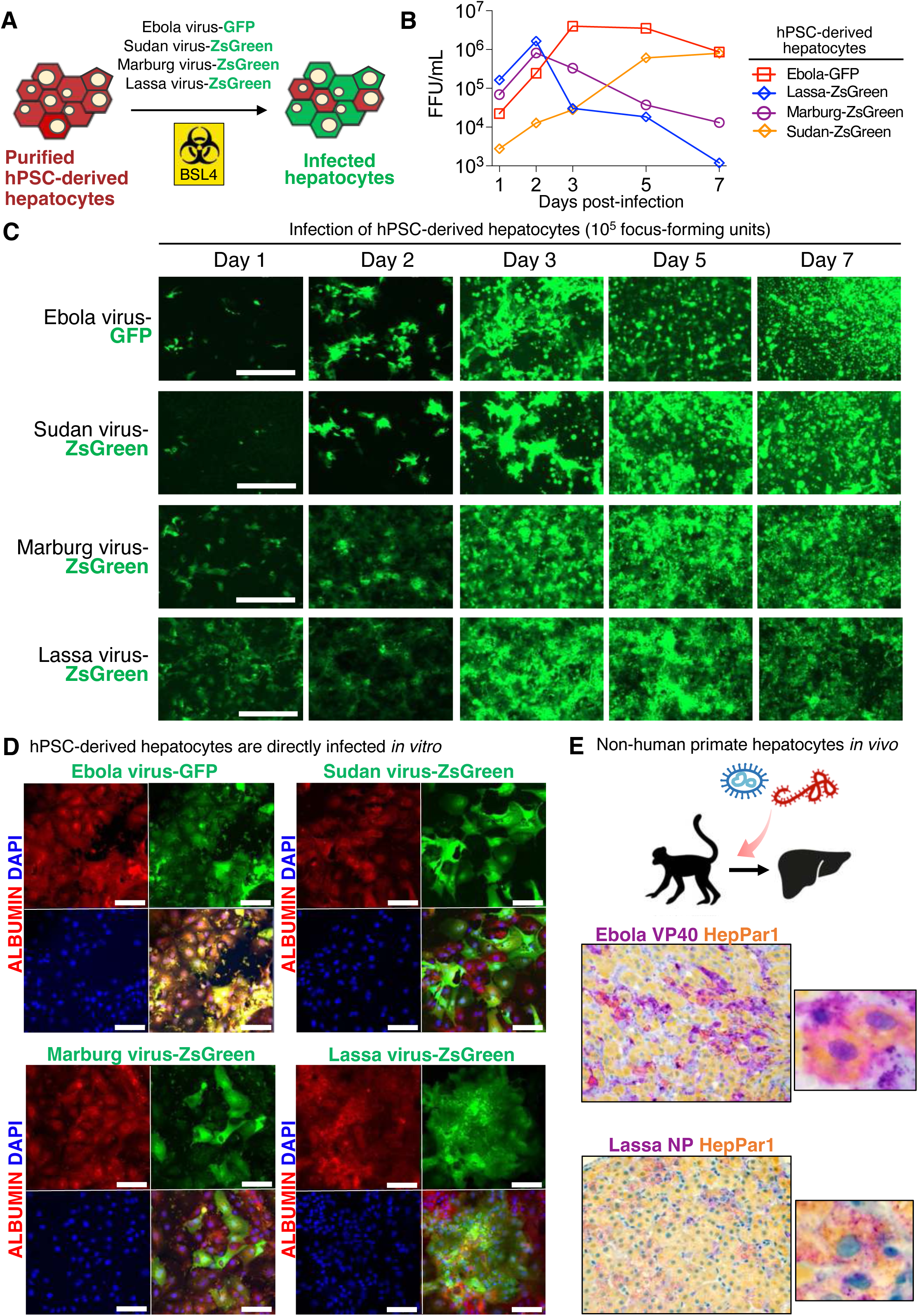
Ebola virus and Lassa virus infect purified hPSC-derived hepatocytes *in vitro* and non-human primate hepatocytes *in vivo*. A) hPSC-derived hepatocytes were purified by metabolic selection, and then cryopreserved, thawed, and cultured for several days before viral inoculation under BSL4 containment. B) hPSC-derived hepatocytes were infected with 10^5^ focus-forming units (FFU) of Ebola virus-GFP, Sudan virus-zsGreen, Marburg virus-zsGreen, or Lassa virus-zsGreen under BSL4 containment. Afterwards, quantification of infectious virus particles in the culture media was performed using the FFU assay, by inoculating Vero cells with hepatocyte-conditioned media. C) Timecourse imaging of hPSC-derived hepatocytes (purified by metabolic selection) infected by 10^5^ FFU of Ebola virus-GFP, Sudan virus-ZsGreen, Marburg virus-ZsGreen, or Lassa virus-ZsGreen. Scale bar, 400μm. D) ALBUMIN immunostaining of hPSC-derived hepatocytes infected with Ebola virus-GFP, Sudan virus-zsGreen, Marburg virus-zsGreen, or Lassa virus-zsGreen for 3 days. Scale bar, 100μm. E) Rhesus or cynomolgus macaques were infected with Ebola or Lassa viruses under BSL4 containment^152, 153^. Liver tissues were immunolabeled for respective viral antigens (purple) and hepatocyte marker HepPar1/CPS1 (yellow), with colocalization of antigens appearing orange or red.

Next, we molecularly confirmed that these Risk Group 4 viruses infect hepatocytes *in vivo*, using archived liver tissues from rhesus or cynomolgus macaques that had been previously infected by Ebola or Lassa virus^152, 153^. Immunohistochemistry revealed colocalization of hepatocyte marker HepPar1/CPS1^154^ and viral antigens within hepatocytes *in vivo* (**Fig. 6E, Fig. S6B**), expanding on past morphologic observations that these viruses infect liver cells *in vivo*^19-21, 23-30^.

### hPSC-derived hepatocytes responded to Ebola and Lassa viruses with divergent effects

What are the effects of Ebola and Lassa viruses on hepatocytes? Although arenaviruses and filoviruses both infect the liver and cause viral hemorrhagic fevers, there are significant differences in disease: Lassa virus (an arenavirus) has a ∼1% fatality rate, whereas filoviruses (e.g., Ebola, Sudan, and Marburg viruses) cause extremely high fatality rates ranging from ∼44-81%^7, 14^. To our knowledge, the effects of arenaviruses and filoviruses on human cells have yet to be compared side-by-side in the same experimental system. In subsequent experiments, we focused on comparing the effects of wild-type (i.e., non-recombinant) Ebola virus (a filovirus) and Lassa virus (an arenavirus) on hPSC-derived hepatocytes. Originally, our Ebola and Lassa virus stocks were titrated on Vero cells. The concentrations of viral stocks measured from Vero cells does not translate 1:1 when applied to hepatocytes. To arrive at equivalent numbers of infected hepatocytes after Lassa or Ebola infection, viral stocks were concentrated, purified, and titrated on hepatocytes (**Fig. S6D**). Of note, hPSC-derived hepatocytes were differentially susceptible to these viruses compared to transformed monkey Vero cells ordinarily used to propagate and titrate these viruses, underscoring the importance of using physiologically-relevant cell-types for viral titration and multiplicity of infection (MOI) calculations (**Fig. S6D**).

We infected purified hPSC-derived hepatocytes with high titers of purified authentic Ebola virus (Mayinga isolate) or Lassa virus (Josiah isolate) to synchronously infect >95% of hepatocytes (MOI ≍ 4) and analyzed transcriptional responses at 6 timepoints (6 hours and 1, 2, 3, 5, and 7 days post-infection). As a control, we also infected cells with Sendai virus (Cantell strain), a non-pathogenic paramyxovirus that stimulates interferon signaling ^155^. Intracellular and extracellular levels of Lassa virus peaked at 1 day post-infection (**Figs. 7A-B**). Intracellular and extracellular Ebola virus levels peaked at 2 and 3 days post-infection, respectively (**Figs. 7A-B**). Ebola virus replication was extensive: it occupied 48.2% of the hepatocyte transcriptome by 2 days post-infection as quantified by RNA-seq (**Fig. 7C**).

**Fig. 7.**
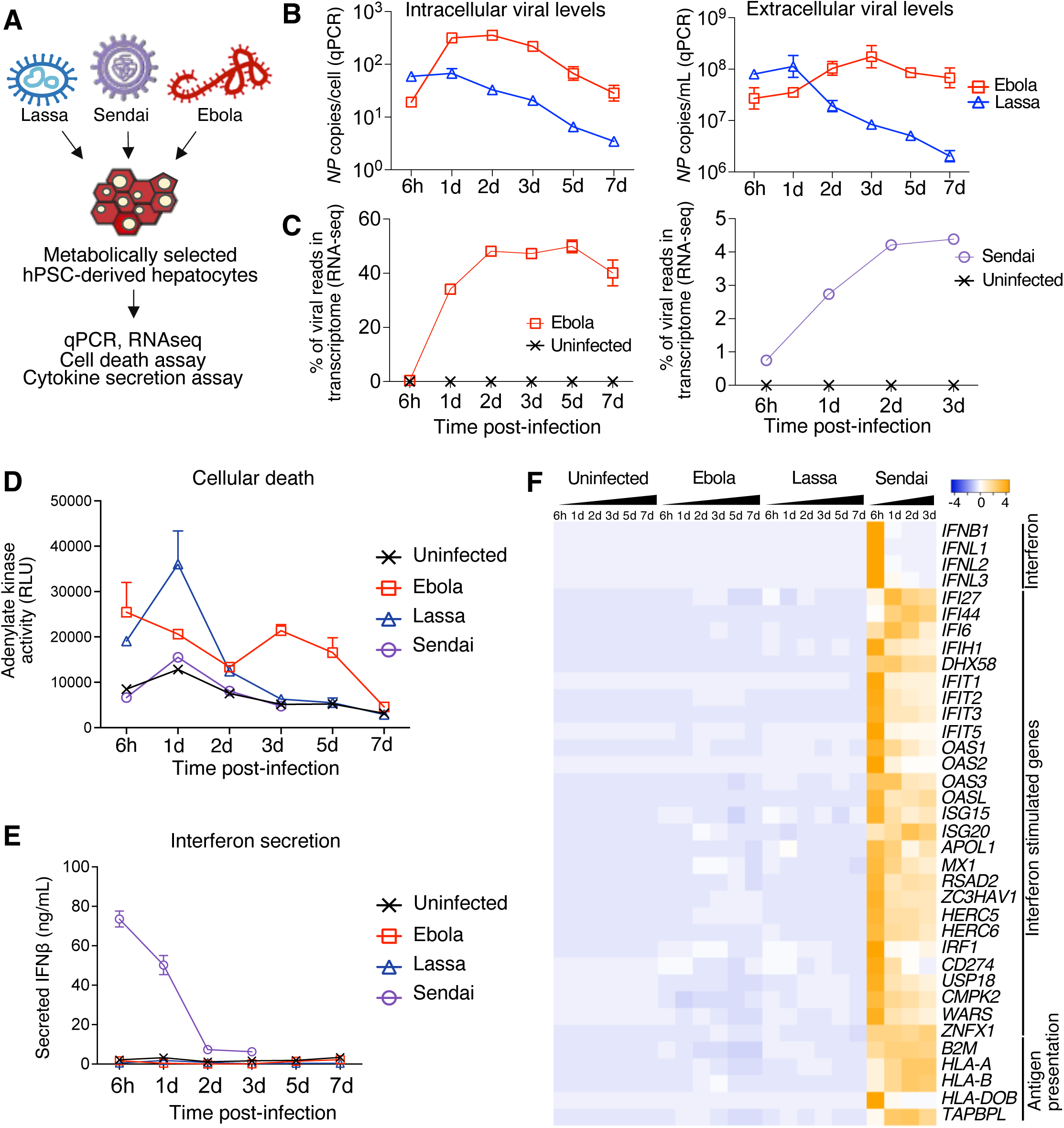
Ebola virus and Lassa virus fail to induce interferon signaling in purified hPSC-derived hepatocytes. A) Metabolically-selected hPSC-derived hepatocytes were infected with wild-type Ebola virus (Mayinga isolate), wild-type Lassa virus (Josiah isolate), or Sendai virus (Cantell strain). High titers of Ebola or Lassa virus were used to synchronously infect most cells (MOI ≍ 4). B) Intracellular levels of Ebola and Lassa viral RNA after inoculating H1 hPSC-derived, metabolically-selected hepatocytes with Ebola or Lassa virus, as quantified by qPCR of the cell monolayer (left; after removing the culture media) or the supernatant (right). Viral *NP* gene copy numbers were respectively normalized to the number of hepatocytes plated for the experiment (left), or volume of the cell culture media (right). C) Percentage of viral reads in the transcriptome of hPSC-derived hepatocytes infected by Ebola or Sendai virus, as determined by RNA-seq. Lassa virus reads were not quantified, as Lassa mRNAs are not polyadenylated and thus were not expected to be captured by our RNA-seq strategy. Error bars denote standard deviations. D) Quantification of cell death of hPSC-derived hepatocytes, either infected by Ebola, Lassa, or Sendai virus, or uninfected. An adenylate kinase assay was performed on the culture media. Adenylate kinase is an intracellular enzyme released into the media upon cell death. Error bars denote standard errors. E) IFNβ secretion by uninfected hPSC-derived hepatocytes, or those infected by Ebola, Lassa, or Sendai virus, as quantified using an enzyme-linked immunosorbent assay (ELISA). Error bars denote standard errors. F) Bulk population RNA-seq of uninfected hPSC-derived hepatocytes, or those infected by Ebola, Lassa, or Sendai virus, showing expression of interferon-stimulated genes^144^. Heatmap shows selected genes.

Ebola virus progressively killed hPSC-derived hepatocytes, as quantified by release of intracellular adenylate kinase into the culture media (**Fig. 7D**, **Fig. S7A**). By contrast, Lassa virus led to a transient wave of cell death at 1 day post-infection (**Fig. 7D**)—coincident with the highest levels of Lassa virus (**Fig. 7B**)—before subsequently re-normalizing.

Ebola and Lassa virus were remarkably effective at suppressing innate immunity in hepatocytes: despite high viral loads, there was almost undetectable secretion of IFNβ protein (**Fig. 7E**) or transcriptional induction of interferons, interferon target genes^144^, or antigen presentation genes (**Fig. 7F**). This was notable, revealing the progressive destruction of Ebola virus-infected hepatocytes (**Fig. 7D**) was not accompanied by interferon production or responses. As a positive control, Sendai virus induced massive interferon secretion and signaling (**Figs. 7E-F**, **Fig. S7B**), indicating that hPSC-derived hepatocytes are competent to produce interferon, but that Ebola and Lassa virus potently evade innate immune detection^14, 156-162^.

If Ebola and Lassa virus do not activate the interferon pathway, what are their transcriptional effects on hepatocytes? The effects of Lassa virus infection were transient and more limited (2 days post-infection, 110 genes upregulated ≥3 fold); they were associated with GO terms such as complement activation, proteolysis, cytolysis, and inflammation (**Figs. 8A-C**), but subsequently largely renormalized within the next few days (by 5 days post-infection, 21 genes upregulated ≥3 fold). By contrast, Ebola virus infection significantly induced the expression of 1475 and 2202 genes after 2 or 5 days, respectively (**Figs. 8A, C**). We found that Ebola virus and Lassa virus up- or downregulated largely different sets of genes (**Figs. 8B, C**).

**Fig. 8.**
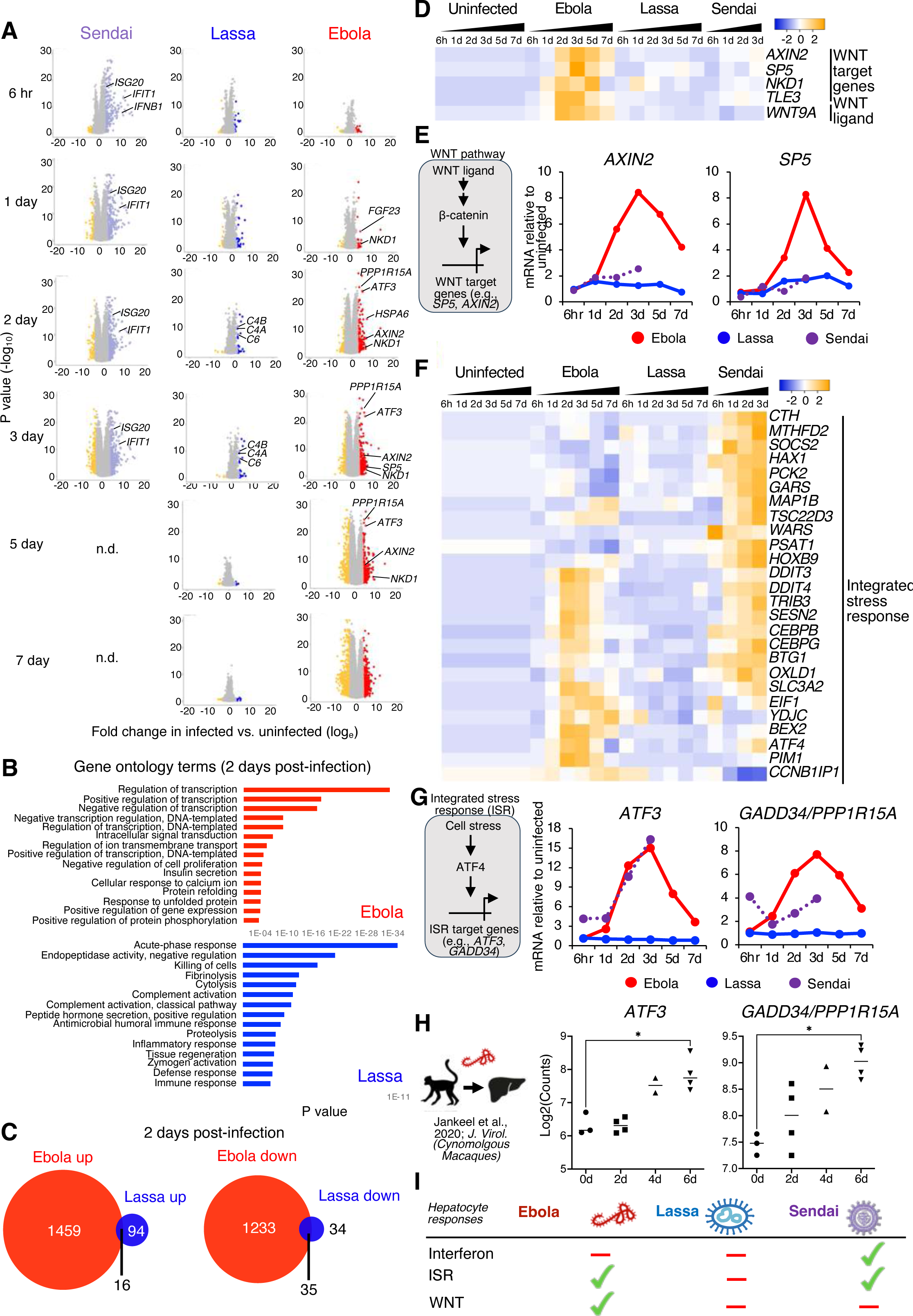
Ebola virus infection activates WNT and ISR pathways in purified hPSC-derived hepatocytes, whereas Lassa virus does not. A) Volcano plots of differentially-expressed genes in Ebola, Lassa, or Sendai virus-infected hepatocytes, relative to mock-infected controls at the same timepoints. In each volcano plot, genes on the left vs. right represent those downregulated or upregulated by viral infection, respectively. n.d. = not determined. B) RNA-seq comparison of Ebola or Lassa virus-infected hPSC-derived hepatocytes vs. mock-infected hepatocytes to determine gene ontology (GO) terms associated with Ebola or Lassa virus infection. C) Venn diagram of genes differentially or commonly up- or downregulated by Ebola and Lassa virus 2 days post-infection of purified hPSC-derived hepatocytes. D-G) Bulk population RNA-seq of purified hPSC-derived hepatocytes, either infected by Ebola, Lassa, and Sendai virus, or uninfected. H) Bulk population RNA-seq of primate liver tissue infected by Ebola over a time course of 0, 2, 4, and 6 days^34^. I) Summary of purified hPSC-derived hepatocyte responses to viral infection.

Remarkably, Ebola virus transcriptionally activated the WNT and integrated stress response (ISR) pathways in hepatocytes, whereas Lassa virus did not. First, Ebola virus induced the expression of WNT ligands (e.g., *WNT9A*) and WNT target genes (e.g., *AXIN2*, *SP5*, and *NKD1*)^163, 164^ (**Figs. 8D-E**). Liver injury rapidly activates WNT signaling to drive regeneration *in vivo*^165^, suggesting that dying, Ebola virus-infected hPSC-derived hepatocytes may engage a pro-regenerative WNT program to compensate for cell death. Second, while Ebola virus did not detectably activate interferon signaling, we found that it instead strongly induced ISR signaling (**Figs. 8F-G**, **Fig. S7C**). ISR is activated by cellular stresses, including viral infection and endoplasmic reticulum (ER) stress, and induces translational shutdown to blunt viral replication^166, 167^. Within 1 day, Ebola virus-infected hepatocytes upregulated a suite of ISR pathway target genes (e.g., *DDIT3/CHOP*, *DDIT4*, *PIM1*, *EIF1*, *CCNB1IP1*, *GADD34/PPP1R15A*), including *ATF3*, a key transcriptional activator of ISR signaling (**Figs. 8F-G, Fig. S7C**)^168, 169^. Consistent with this *in vitro* finding, our analysis of published RNA-seq datasets of Ebola-infected non-human primates^34, 170^—which represent the gold standard model for Ebola virus disease—likewise revealed *ATF3* and *PPP1R15A* were upregulated in the Ebola-infected primate liver (**Fig. 8H, Fig. S7E**).

By contrast, ISR and WNT genes were not induced by Lassa virus *in vitro*, suggesting this is not a generic hepatocyte response to viral infection (**Figs. 8F-G, Fig. S7C**). Sendai virus infection also induced certain ISR genes, some of which were different (*GARS*, *HAX1*, and *WARS*) than those induced by Ebola (*CCNBP1IP1*, *YDJC*, and *PIM1*), potentially reflecting activation of different ISR branches or differing levels of ISR activation (**Figs. 8F-G, Fig. S7C**). Finally, Ebola virus and Sendai virus, but not Lassa virus, reduced expression of various liver function genes, including *FGB*, *ALB*, *TTR*, *APOE*, and *ARG1* (**Fig. S7D**). Taken together, there are starkly different effects of Ebola virus and Lassa virus on purified human hepatocytes, especially with regard to cell death and activation of WNT and ISR pathways (**Fig. 8I**).

## DISCUSSION

Here we developed a new approach to create pure human hepatocytes *in vitro* from hPSCs, and we applied these purified hPSC-derived hepatocytes to discover how Ebola virus and Lassa virus affect hepatocytes. First, we differentiated hPSCs into ∼80% pure hepatocytes, and pioneered a “metabolic selection” approach—withholding glucose, glutamine, and pyruvate—to subsequently deplete non-liver cells with high speed and specificity, providing a convenient method to purify hepatocytes from heterogeneous cell populations. Purified hPSC-derived hepatocytes were transcriptionally and functionally more akin to primary hepatocytes than liver cancer cell lines prevalently used in BSL4 virology. While Ebola and Lassa virus extensively infected hepatocytes *in vivo* and *in vitro*, we found that they led to starkly different effects. Ebola infection activated the WNT and ISR pathways in human hepatocytes, whereas Lassa did not. To our knowledge, this is the first time that different Risk Group 4 viral families (e.g., filoviruses vs. arenaviruses) have been directly compared in the same human experimental system, thus representing a step forward for comparative virology. In sum, purified hPSC-derived hepatocytes provide an abundant, experimentally accessible, and physiologically relevant model system for BSL4 virology. In turn, this will accelerate studies of the basic biology of these viruses and future therapeutic screens.

### A roadmap for human liver differentiation

We comprehensively profiled stepwise changes in gene expression, chromatin accessibility, and cellular diversity of each step of hPSC differentiation into hepatocytes, using scRNAseq and OmniATACseq. This provides a rich resource (https://anglab.shinyapps.io/liverscrna/) to discover new markers and regulators of human liver differentiation. This resource also encompasses freshly-isolated and cultured adult hepatocytes and liver cancer cell lines (HepG2 and HuH-7), allowing for unbiased benchmarking of different liver cell culture models.

At each stage of differentiation, we provided lineage-specifying cues to generate the cell-type of interest while, of equal importance, blocking the signals that generated unwanted cell-types. scRNAseq revealed that this two-pronged approach rapidly and efficiently differentiated hPSCs into primitive streak, definitive endoderm, posterior foregut, and liver bud progenitors within 1, 2, 3, and 6 days of differentiation. Cell populations were mostly uniform at each of these steps, with little evidence of unwanted cell-types arising and reiterating the high precision of early differentiation (**Fig. S1D**).

By day 18 of differentiation, ∼80% pure hepatocytes emerged, although we discovered they were commingled with intestinal goblet and neuroendocrine cells. We thus document the identity of non-liver cells that arise alongside hepatocytes during hPSC differentiation. Indeed, the liver and intestine are adjacent endodermal organs^107^, and it is thus plausible for intestinal cells to erroneously appear during liver differentiation *in vitro*^171^. The emergence of these non-liver cells necessitates approaches to exclusively purify hepatocytes, as heterogeneous cell populations pose challenges for various applications, including virology, drug screening, and regenerative medicine.

### Metabolic selection: a new approach to purify hepatocytes

Purifying desired cell-types from heterogeneous cell populations remains a significant challenge for stem cell biology and regenerative medicine^54^. To purify hepatocytes, we developed “metabolic selection,” a new approach to rapidly kill non-liver cells simply by withholding certain essential nutrients from the culture medium. This builds on the emerging concept that different cell-types require specific nutrients to survive^82-87^. Indeed, hepatocytes are one of the few cell-types capable of converting glucose into glycogen^88-91^, and can therefore rely on glycogen during glucose deprivation. Withholding glucose, pyruvate, and glutamine in HepSelect media for 1-3 days destroyed non-liver cells *in vitro*, while hepatocytes survived. Metabolic selection generated essentially pure hPSC-derived hepatocytes; scRNAseq could not overtly detect surviving intestinal or other cell-types. Importantly, metabolic selection purifies hepatocytes based on their metabolic functionality instead of surface marker expression^92^. Metabolic selection thus provides a simple, scalable, and inexpensive method to purify hepatocytes.

Moreover, the combinatorial withdrawal of nutrients was critical to purify hepatocytes. Glucose removal was insufficient to eliminate intestinal cells (**Fig. S3C**). A previous study showed glucose removal was also insufficient to destroy hPSCs because they can utilize either glutamine or arginine to survive in glucose-depleted media^119, 120^. We found that simultaneously withholding glucose, pyruvate, and glutamine was critical to purify hepatocytes and eliminate non-liver cells. More broadly, metabolic selection might serve to purify other cell-types, including neurons, adipocytes, kidney, and skeletal muscle cells^172-174^ that also synthesize glucose *de novo* or rely on non-glucose sources to produce energy.

### hPSC-derived hepatocytes provide an enhanced model system for BSL4 virology

The unique constraints of BSL4 experimentation^37^ have long posed a formidable challenge to understanding the mechanistic effects of Risk Group 4 viruses on human hepatocytes. Thus far, liver cancer cell lines such as HepG2 and HuH-7 have been extensively used in BSL4 virology^38-43^, but cannot produce interferon and lack many hepatocyte features. hPSC-derived cell-types offer key advantages for BSL4 virology, as they can be produced in large numbers, yet are chromosomally normal, resemble their *in vivo* counterparts, and are not oncogenically transformed^48, 175, 176^. However, one drawback of hPSC-derived cellular models is cellular heterogeneity, as hPSC differentiation often yields complex mixtures of multiple cell-types^177^. Indeed, a pioneering study found that Ebola virus could infect differentiated hPSC populations that contained ∼25% ALBUMIN+ hepatocytes but predominately contained non-liver cells^48^. However, it was unclear whether hepatocytes and/or non-liver cells were infected, and whether any of the observed cellular responses could be attributed to non-liver cells within the population^48^. Our ability to create purified human hepatocytes from hPSCs—which we showed can be extensively infected by Ebola virus, Sudan virus, Marburg virus, and Lassa virus— provides a more precise platform to study the detailed cellular effects of these Risk Group 4 viruses.

### Effects of Ebola virus on human hepatocytes

Multiple Risk Group 4 viruses, including Ebola virus and Lassa virus, have a predilection to infect hepatocytes. While Ebola virus also infects other tissues in the body, liver infection has been implicated in hepatocyte death, coagulopathies, and liver injury that often accompanies end-stage disease^6, 21, 178, 179^. Hepatocytes secrete voluminous amounts of proteins (e.g., carrier and coagulation proteins) into the bloodstream, and much of their cell body is devoted to protein synthesis and secretion^180^. We find that Ebola virus occupies ∼50% of the hepatocyte transcriptome for up to a week, during which high levels of infectious Ebola virus particles are continuously released into the media. While immune cells are initial targets of Ebola virus *in vivo*, it is possible that once Ebola virus reaches the liver, it transforms hepatocytes into efficient virus production factories to achieve systemic viral dissemination and to overwhelm the host. Indeed, we and others previously discovered that the early extent of liver replication correlates with Ebola virus disease severity in humanized mice^181, 182^. *In vivo*, Ebola virus reaches extraordinary bloodborne levels within several days^179^, which could partly reflect massive viral secretion by hepatocytes in the absence of effective immunological restraint of viral replication.

Ebola virus fails to trigger interferon and inflammatory cytokine expression by infected hepatocytes, despite massive viral loads (∼50% of the transcriptome) for up to a week. High-level viral production in the virtual absence of interferon production may allow Ebola virus to stealthily replicate in, and be released from, infected hepatocytes without initially engaging immune defenses that might curb viral replication. Such interferon suppression is not a feature of all liver-tropic viruses: by contrast, hepatitis C virus potently induces interferon in hepatocytes^183^. Additionally, Ebola virus-infected hepatocytes do not express inflammatory cytokines (e.g., CXCL1, CXCL10, and CXCL12) that recruit T cells, NK cells, and myeloid cells^184^, which are instead strongly expressed by Sendai-virus infected hepatocytes. Initial impairment of immune cell recruitment into the liver may help explain the delayed immune responses observed in Ebola virus disease^7, 14^. Our findings that Ebola virus-infected hepatocytes minimally produce interferon or inflammatory cytokines may also help explain a longstanding curiosity in the field: there can be extensive hepatocyte necrosis, yet minimal inflammation, in Ebola virus-infected livers^21^.

The conspicuous absence of interferon or inflammatory cytokine production after Ebola virus infection does not reflect a fundamental deficiency of our model system. We show that hPSC-derived hepatocytes are competent to secrete such cytokines in response to Sendai virus or poly(I:C) treatment, unlike liver cancer cell lines^46-48^. However, our findings contrast with a recent study that infected a ∼25% pure population of hPSC-derived hepatocytes with Ebola virus, and described interferon mRNA upregulation, although interferon protein levels were not assessed^48^. The divergent results between our two studies could be attributed to unidentified, non-liver cells present within the culture, emphasizing the importance of generating pure hepatocytes. Overall, our results are generally consistent with autopsies that revealed surprisingly minimal inflammation in Ebola virus-infected livers^21^.

While Ebola virus did not trigger interferon production, we found that hepatocytes nevertheless responded in two unexpected ways: activating the WNT and ISR pathways. First, we find that Ebola virus infection induces WNT signaling in hPSC-derived hepatocytes. Liver injury induces WNT signaling, thus triggering hepatocyte proliferation and regeneration^165^. Our results could help explain human autopsy results that there is abundant hepatocyte proliferation in Ebola virus-infected livers, thought to reflect an ongoing regenerative response to cell death^4^. Second, the ISR pathway can be triggered by multiple types of cellular stress, including double-stranded RNA (sensed by PKR) and ER stress, perhaps elicited by overproduction of viral proteins that cannot be correctly folded (sensed by PERK) ^166, 167^. The upstream drivers and downstream consequences of ISR pathway activation in Ebola virus infection are presently unknown. Classically, ISR activation leads to translational shutdown to curtail viral replication^166, 167^, which could present a cell-intrinsic mechanism for hepatocytes to reduce viral protein production despite failed interferon pathway activation. Additionally, excessive ISR signaling can eventually lead to cell death^166, 167^, providing a potential mechanism for Ebola virus-induced hepatocyte death that we observed.

### Differences in how Ebola and Lassa viruses affect human hepatocytes

To our knowledge, this is the first study comparing the cellular effects of Risk Group 4 viruses from two different families—Ebola virus (a filovirus) vs. Lassa virus (an arenavirus)—side-by-side in the same human experimental system. Comparative virology, whereby the effects of different viruses are compared and contrasted, is a major pursuit of modern virology. The effects of different filoviruses have been compared^181, 182, 185-190^. However, the effects of a Risk Group 4 filovirus (Ebola virus) vs. an arenavirus (Lassa virus) on human cells remain an open question^14, 17^.

Curiously, even after infection with a high viral dose (MOI ≍ 4), Lassa virus induced modest and transient transcriptional changes in hepatocytes. Despite infecting up to 500,000 individuals every year in Africa, ∼80% of Lassa cases are mild or asymptomatic^14, 191^. However, we find that despite high viral loads, Lassa is similar to Ebola virus in that it suppressed production of both interferon and inflammatory cytokines from infected hepatocytes. This may facilitate initial Lassa virus infection of hepatocytes, consistent with results from autopsies^20, 24-26^, while impairing early immune system engagement. However, we find that Lassa virus does not activate the WNT or ISR pathways, a key difference from Ebola. Our findings are altogether consistent with a hallmark autopsy study that although Lassa virus infects the human liver, its effects on the liver are generally more mild^20^.

More broadly speaking, we demonstrate that purified hPSC-derived hepatocytes provide a powerful experimental system to generate and test mechanistic hypotheses regarding Risk Group 4 viruses, which would otherwise prove challenging in other model systems given the experimental constraints of BSL4 containment. hPSC-derived hepatocytes also constitute a platform to study and compare additional Risk Group 4 viruses, including nairoviruses (e.g., Crimean-Congo hemorrhagic fever virus^192-195^) and New World arenaviruses (e.g., Junin virus ^196^) that infect human hepatocytes *in vivo* but whose cellular effects remain obscure.

## ACKNOWLEDGMENTS

This work is dedicated to the memory of Siew Lan Soh. We also thank Andrew Elefanty, Ed Stanley, and Elizabeth Ng for *MIXL1-GFP* hPSCs; Emily Rae Dwyer, Faith-Masong Njunkeng, Jiayi Wu, Meng Zhao, Katrin Svensson, Alana Nguyen, Jun Jiang, and Antson Tan for contributing to experiments and analyses; Kyle Cromer, Phil Beachy, Roel Nusse, Hiromitsu Nakauchi, Aaron Kershner, Massimo Nichane, Siva Vijayakumar, Aijaz Ahmed, Natalie Torok, Wendy Wenderski, Xiaochen Xiong, Jiyun Choi, Reuben Saunders, Aditya Anand, Ren Ee Chee, and Yoshihiro Kawaoka for reagents and advice; and Trenton Bushmaker and Kyle Rosenke for sharing non-human primate tissues. Infrastructure support was provided by the Stanford Institute for Stem Cell Biology & Regenerative Medicine, Stanford Stem Cell FACS Core Facility, Stanford Diabetes Genomics Analysis Core Facility, Valerie Park, Liying Ou, Laura Dunkin-Hubby, Catherine Carswell-Crumpton, Joe Olage Pasillas III, and Cheng Pan. This work was supported by the California Institute for Regenerative Medicine (DISC2-10679), Stanford Maternal and Child Health Research Institute, U.S. National Institutes of Health (NIH Director’s Early Independence Award DP5OD024558 and P30DK116074), JDRF Northern California Center of Excellence, Stanford Beckman and Ludwig Centers, Robert Koch Institute intramural funds, and the Anonymous, Fickel, Gilbert, and Weintz families. K.R.-H. is an HHMI Investigator. K.M.L. is a Human Frontier Science Program Young Investigator (RGY0069/2019), Packard Foundation Fellow, Pew Scholar, Baxter Foundation Faculty Scholar, and The Anthony DiGenova Endowed Faculty Scholar. L.T.A. is a Siebel Investigator and an Additional Ventures Career Development Awardee.

## AUTHOR CONTRIBUTIONS

K.J.L., N.M.Q.P., Manali B., K.M.L., and L.T.A. generated and characterized hPSC-derived hepatocytes. J.B.P., Marcel B., and A.L. performed BSL4 experiments. L.T.A., K.J.L., P.W.K., H.Y., and K.M.L. analyzed RNA-seq. S.K.J. and K.R.H. generated PROX1-2A-GFP hPSC reporter line. L.T.A. performed surface marker screening and Omni-ATAC-seq analysis. B.L. and I.L.W. provided advice. With input from K.M.L., L.T.A. and J.B.P. supervised the study.

## DECLARATION OF INTERESTS

Stanford University has filed patent applications related to this work. B.L. was involved with this study when employed by the Genome Institute of Singapore; he is now at Thymmune Therapeutics, which had no role in the present study.

## DATA AND CODE AVAILABILITY

The bulk-population RNA-seq and single-cell RNA-seq datasets generated in this study are available at the NCBI Sequence Read Archive, accession number SUB12011576. Single-cell RNA-seq datasets of hPSC-derived hepatocytes developed in this study can be interactively browsed at a custom web portal: https://anglab.shinyapps.io/liverscrna/. Computational scripts used for the genomic analysis conducted in this study are available on GitHub: https://github.com/LayTengAngLab/Liver-BSL4-viruses.

## SUPPLEMENTAL FIGURE LEGENDS

**Fig. S1.**
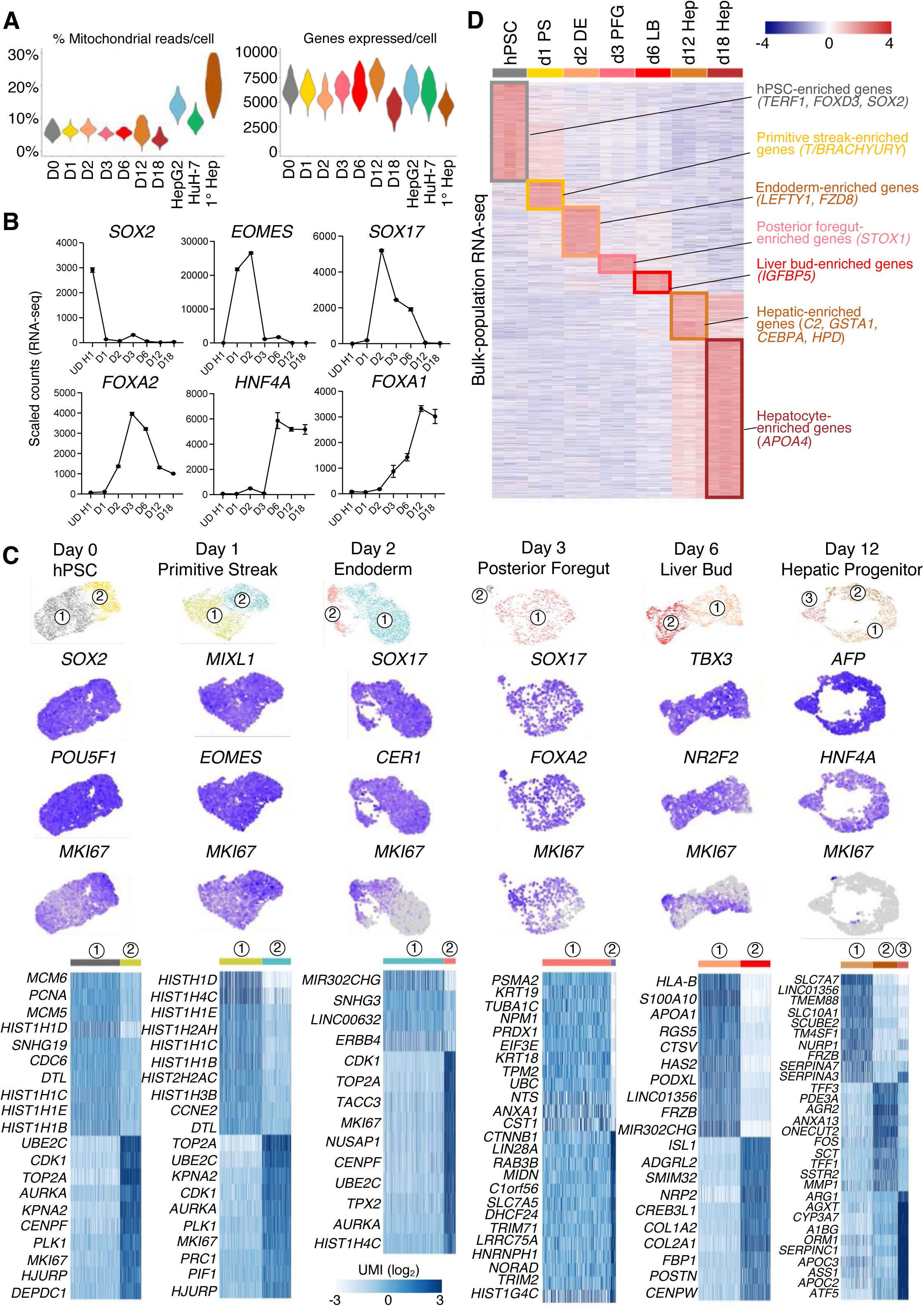
Stepwise changes in gene expression during human liver differentiation from hPSCs, related to Fig. 1. A) Quality control statistics for scRNAseq analysis, showing the number of genes expressed per cell and the percentage of mitochondrial reads over total reads per cell for hPSCs (D0) differentiated into hepatocytes (days 1-18 [d1-d18] of differentiation), HepG2, HuH7, and fresh primary human hepatocytes (1° Hep). B) Bulk-population RNA-seq of H1 hPSCs differentiated into hepatocytes, showing stepwise changes in transcription factor expression. Gene expression quantified in scaled count units. Cell-types profiled include undifferentiated hPSCs (UD), day 1 anteriormost primitive streak, day 2 definitive endoderm, day 3 posterior foregut, day 6 liver bud progenitors, day 12 early hepatocytes, and day 18 hepatocytes. C) scRNAseq of hPSCs (D0) differentiated towards early hepatocytes (days 1-12 [d1-d12] of differentiation) to assess the presence of potential population heterogeneity at each step of differentiation. Louvain clustering^199^ was applied to decompose each cell population into subclusters, and differentially-expressed genes that distinguish these subclusters were identified. Gene expression quantified in log_2_ UMI units. D) Bulk-population RNA-seq of H1 hPSCs, day 1 anteriormost primitive streak, day 2 definitive endoderm, day 3 posterior foregut, day 6 liver bud progenitors, day 12 early hepatocytes, and day 18 hepatocytes, highlighting cell-type-specific gene expression.

**Fig. S2.**
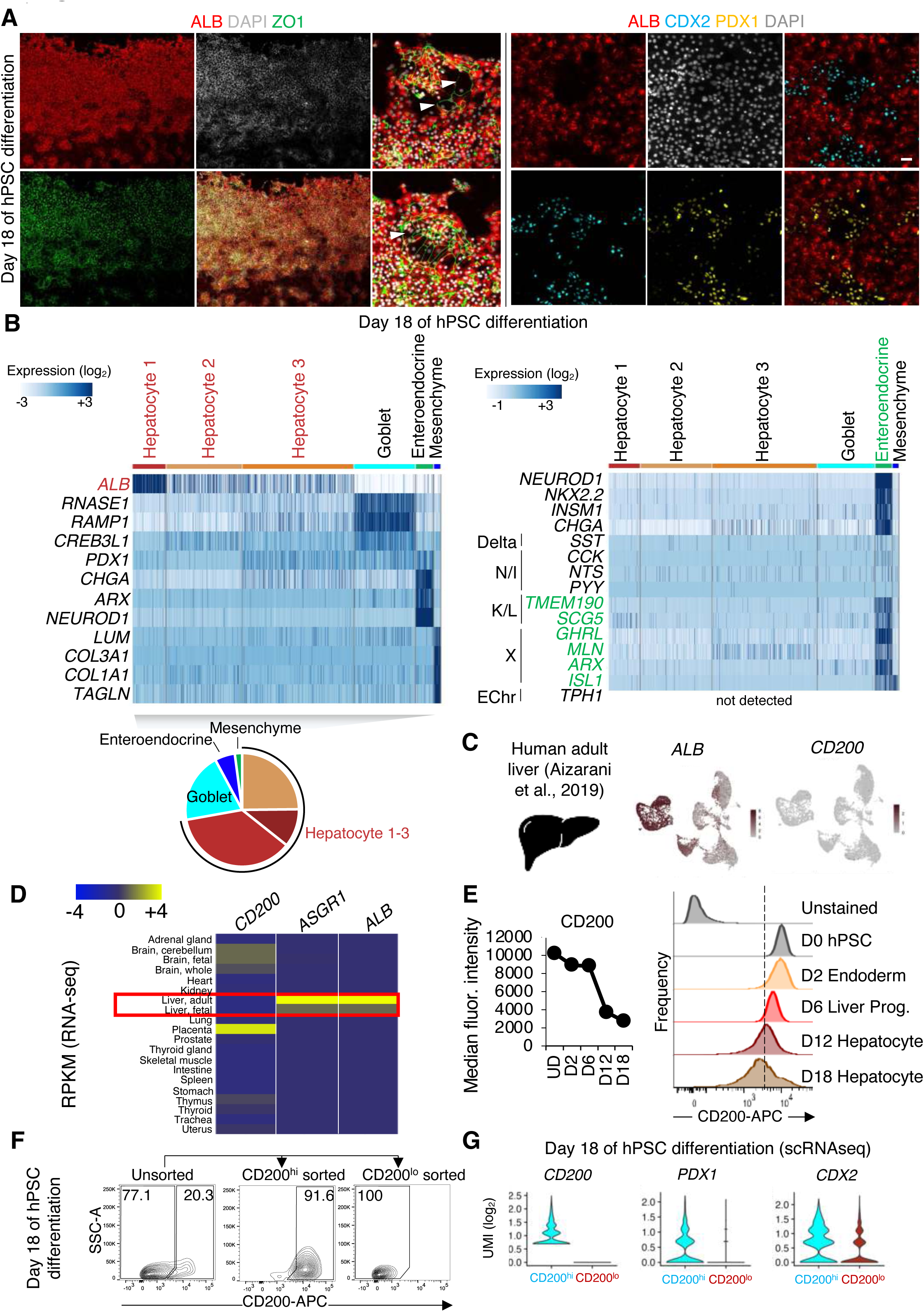
Discovery of non-liver cells generated during liver differentiation from hPSCs, related to Fig. 2. A) ALBUMIN, ZO1, CDX2, PDX1, and DAPI immunostaining of H1 hPSC-derived day 18 hepatocyte-containing populations. Scale bar, 50μm. B) scRNAseq of day 18 H1 hPSC-derived hepatocyte-containing populations, followed by Louvain clustering^199^ to identify multiple cell-types within the population. The entire cell population was harvested without pre-selection. In the heatmap, each column represents gene expression of a single cell, clustered together by “cell-type” clusters (*top*). The proportions of each identified cell-type within the day 18 population are also shown (*bottom*). Three subsets of non-liver cells were discovered: intestinal goblet, intestinal enteroendocrine, and mesenchymal cells. Markers of different intestinal enteroendocrine cell subtypes^111, 112^ were also quantified (*right*), revealing expression of K/L and X enteroendocrine subtype markers. Gene expression quantified in log_2_ UMI units. C) scRNAseq of adult human liver^118^ reveals the absence of *CD200* expression. D) Bulk-population RNA-seq of different human adult tissues^117^ reveals the absence of *CD200* mRNA expression from the liver. E) Flow cytometry of CD200 expression on undifferentiated hPSCs, day 2 definitive endoderm, day 6 liver bud progenitors, day 12 early hepatocytes, and day 18 hepatocytes, showing median fluorescence intensity (*left*) and population expression levels (*right*). F) Flow cytometry of day 18 hPSC-derived hepatocyte-containing populations, showing FACS gates used to sort CD200^hi^ vs. CD200^lo^ populations that were analyzed by qPCR in **Fig. 2f**. G) scRNAseq of hPSC-derived day 18 hepatocyte-containing populations that were computationally binned based on *CD200* mRNA expression into *CD200*^hi^ vs. *CD200*^lo^ subsets.

**Fig. S3.**
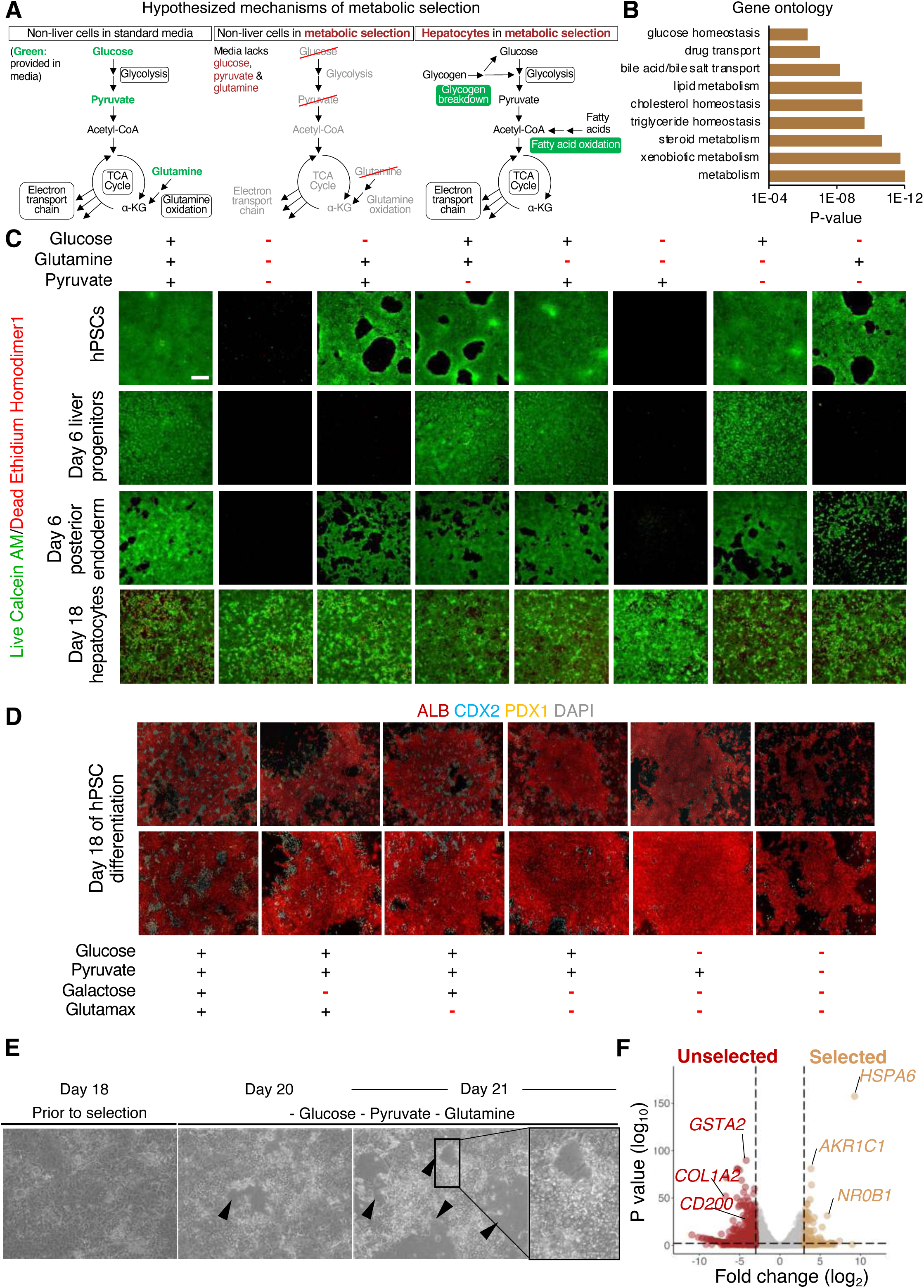
Optimization of metabolic selection to purify hPSC-derived hepatocytes, related to Fig. 3. A) Hypothesized mechanisms of metabolic selection. B) Bulk-population RNA-seq of genes upregulated in day 18 H1 hPSC-derived hepatocytes over undifferentiated hPSCs, followed by gene ontology (GO) analysis of hepatocyte-upregulated genes. This revealed transcriptional upregulation of various metabolic pathways in hepatocytes. C) Cell viability staining of undifferentiated hPSCs, day 6 liver progenitors, day 6 midgut/hindgut (posterior) endoderm, and day 18 hepatocytes cultured in standard vs. metabolic selection media for 1-2 days. Scale bar, 500μm. D) ALBUMIN, CDX2, PDX1, DAPI immunostaining of H1 hPSC-derived day 18 hepatocytes cultured in standard vs. metabolic selection media for 2 additional days. E) Brightfield images of day 18 H1 hPSC-derived hepatocytes that were cultured for 2-3 additional days in metabolic selection media, showing cell death after metabolic selection. F) Bulk-population RNA-seq of day 18 H1 hPSC-derived hepatocytes before, or after culture in metabolic selection media for 1 day to identify genes induced or repressed by metabolic selection.

**Fig. S4.**
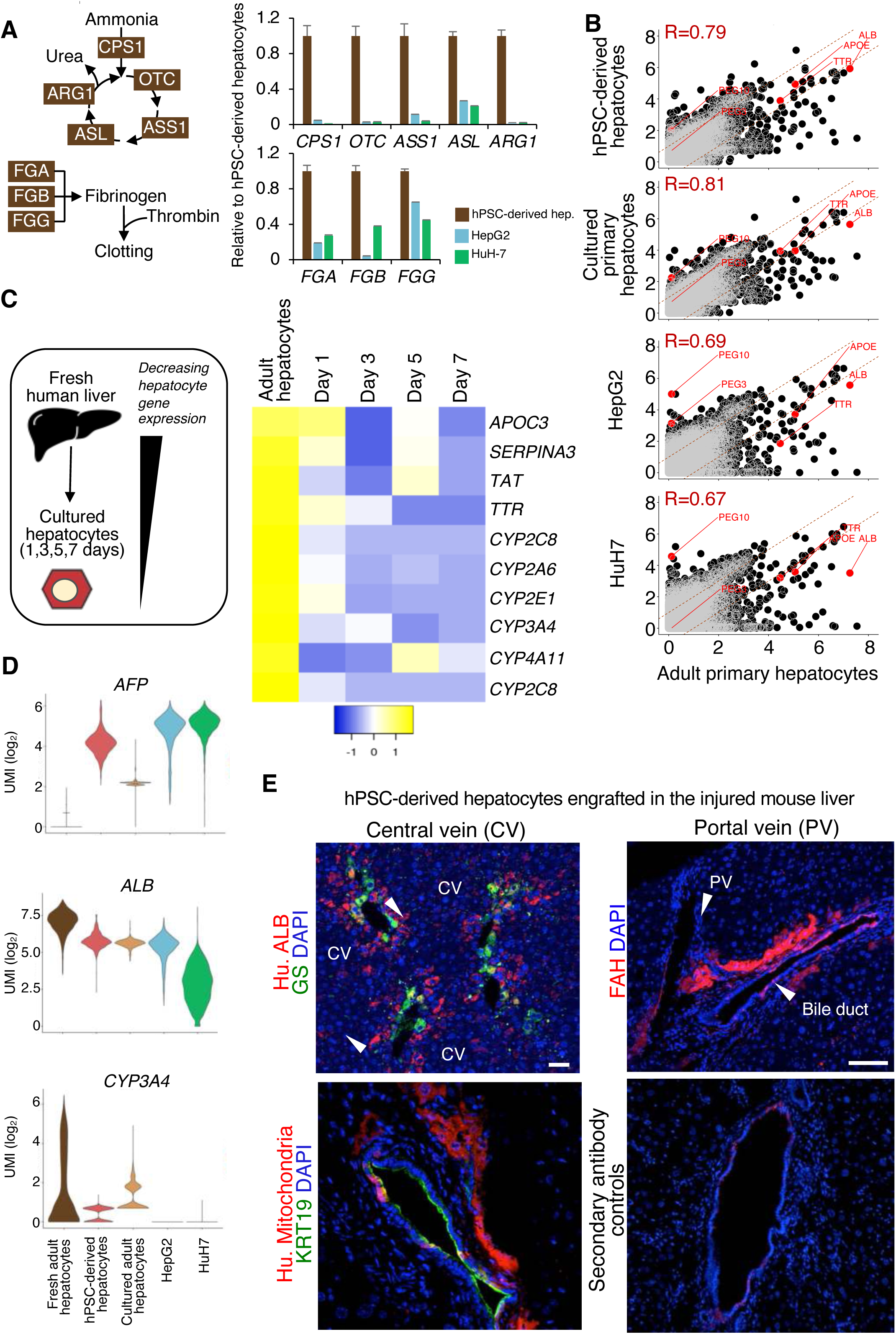
Additional comparisons of metabolically-selected hPSC-derived hepatocytes, adult primary hepatocytes, and liver cancer cells, related to Fig. 4. A) Bulk-population RNA-seq of metabolically-selected hPSC-derived hepatocytes, HepG2, and HuH7 cells. Gene expression normalized to levels found in hPSC-derived hepatocytes. B) Global transcriptome comparison of metabolically-selected hPSC-derived hepatocytes, cultured primary hepatocytes, HepG2, and HuH7 with freshly thawed adult primary hepatocytes. Transcriptome-scale Pearson correlation coefficients between each pair of cell-types are shown. C) Bulk population RNA-seq performed of primary adult human hepatocytes were cultured for 1-7 days in monolayers. D) scRNAseq analysis of metabolically-selected H1 hPSC-derived hepatocytes, liver cancer cell lines (HepG2 and HuH7), and freshly thawed and cultured adult human hepatocytes.

**Fig. S5.**
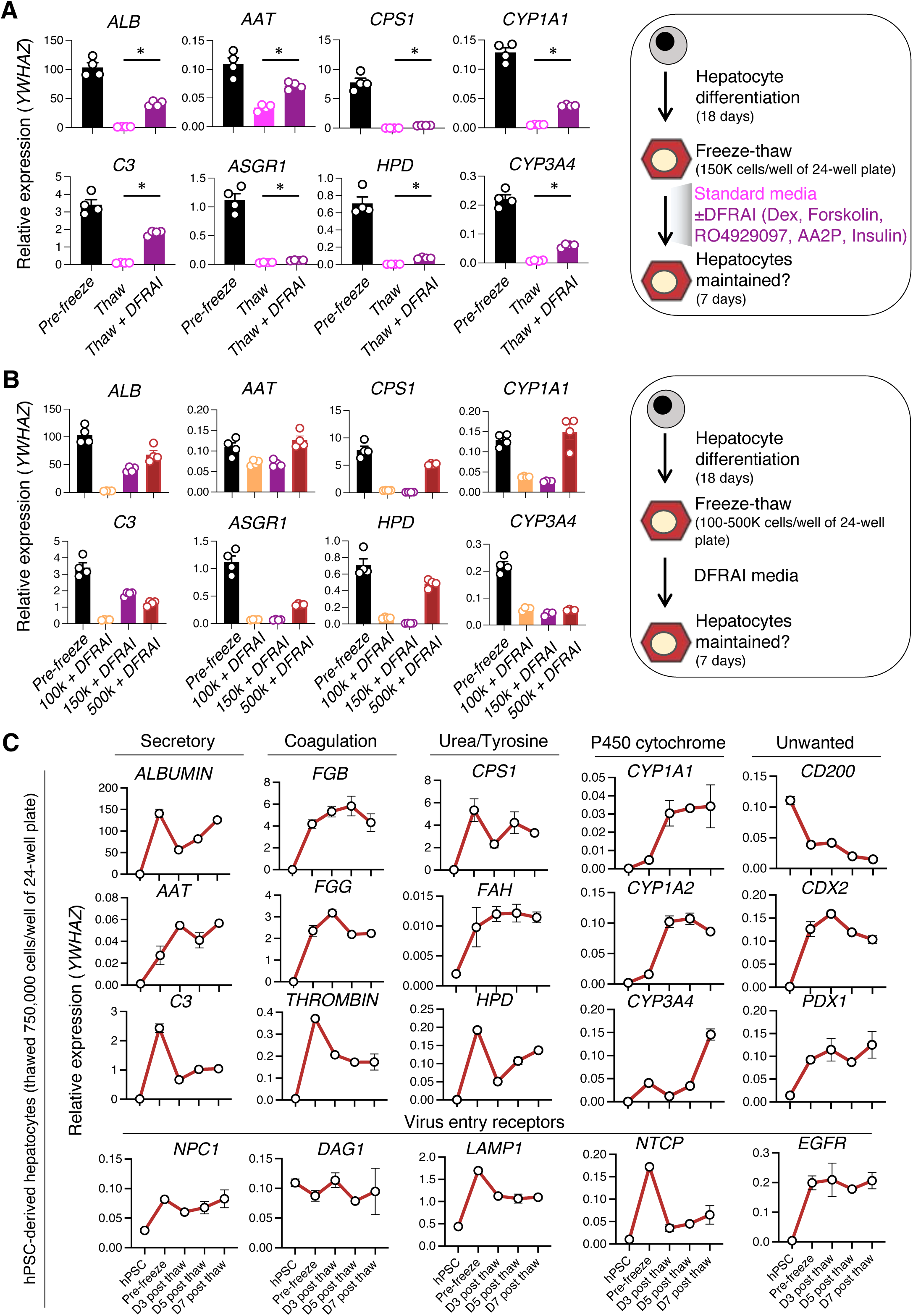
Cryopreservation, thawing, and optimized *in vitro* culture of hPSC-derived hepatocytes, related to Fig. 6. A) qPCR of H1 hPSC-derived day 18 hepatocytes that were cryopreserved, and then thawed at a density of 150,000 cells/well of a 24-well plate either in standard media or media supplemented with hepatocyte-specifying signals (DFRAI: dexamethasone + forskolin + RO4929097 + ascorbic acid-2-phosphate [AA2P] + insulin^69^) for 1 week. Gene expression data normalized to *YWHAZ*, such that *YWHAZ* expression = 1.0. Error bars denote standard errors. B) qPCR of H1 hPSC-derived day 18 hepatocytes that were cryopreserved, then thawed at a density of 100,000-500,000 cells/well of a 24-well plate in DFRAI media for 1 week. Gene expression data normalized to *YWHAZ*, such that *YWHAZ* expression = 1.0. Error bars denote standard errors. C) qPCR of undifferentiated H1 hPSCs, day 18 hPSC-derived hepatocytes before freeze, or hPSC-derived hepatocytes that had been thawed and cultured for 3, 5, or 7 days (D3-7) in DFRAI media at a density of 750,000 cells/well of a 24-well plate. Gene expression data normalized to *YWHAZ*, such that *YWHAZ* expression = 1.0. Error bars denote standard errors.

**Fig. S6.**
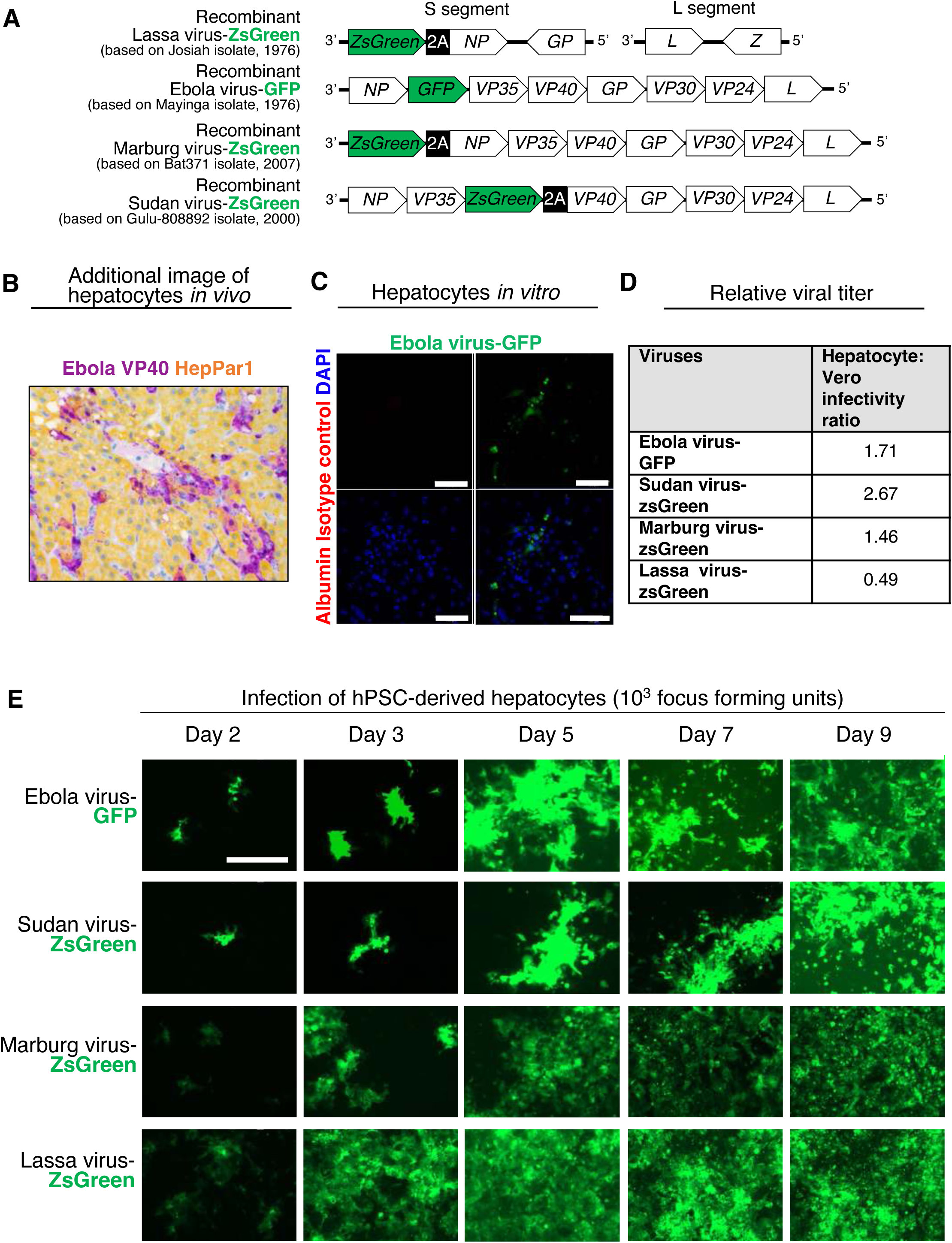
Ebola, Sudan, Marburg, and Lassa virus extensively infect purified populations of hPSC-derived hepatocytes, related to Fig. 6. A) Genome structures of recombinant GFP-expressing Ebola virus^149^, ZsGreen-expressing Sudan virus^150^, ZsGreen-expressing Marburg virus^151^, and ZsGreen-expressing Lassa virus ^43^. B) Additional image of a Ebola virus-infected rhesus macaque liver^153^ immunolabeled for Ebola virus VP40 and hepatocyte marker HepPar1/CPS1. This rhesus macaque was from the same study^153^ shown in **Fig. 6e**. C) Isotype control immunostaining of hPSC-derived hepatocytes infected with Ebola virus-GFP. Scale bar 100μm. Negative control for **Fig. 6d**. D) Relative infectivity of Vero E6 cells vs. H7 hPSC-derived hepatocytes infected with either Ebola virus-GFP, Sudan virus-zsGreen, Marburg virus-zsGreen, or Lassa virus-zsGreen. Both cell-types were inoculated with the same viral dose, and infected cells were quantified using the fluorescent FFU assay. Higher numbers indicate that hPSC-derived hepatocytes were preferentially infected by the virus, relative to Vero cells. E) Timecourse imaging of hPSC-derived hepatocytes (purified by metabolic selection) infected by 10^3^ FFU of Ebola virus-GFP, Sudan virus-ZsGreen, Marburg virus-ZsGreen, or Lassa virus-ZsGreen. Scale bar, 400μm.

**Fig. S7.**
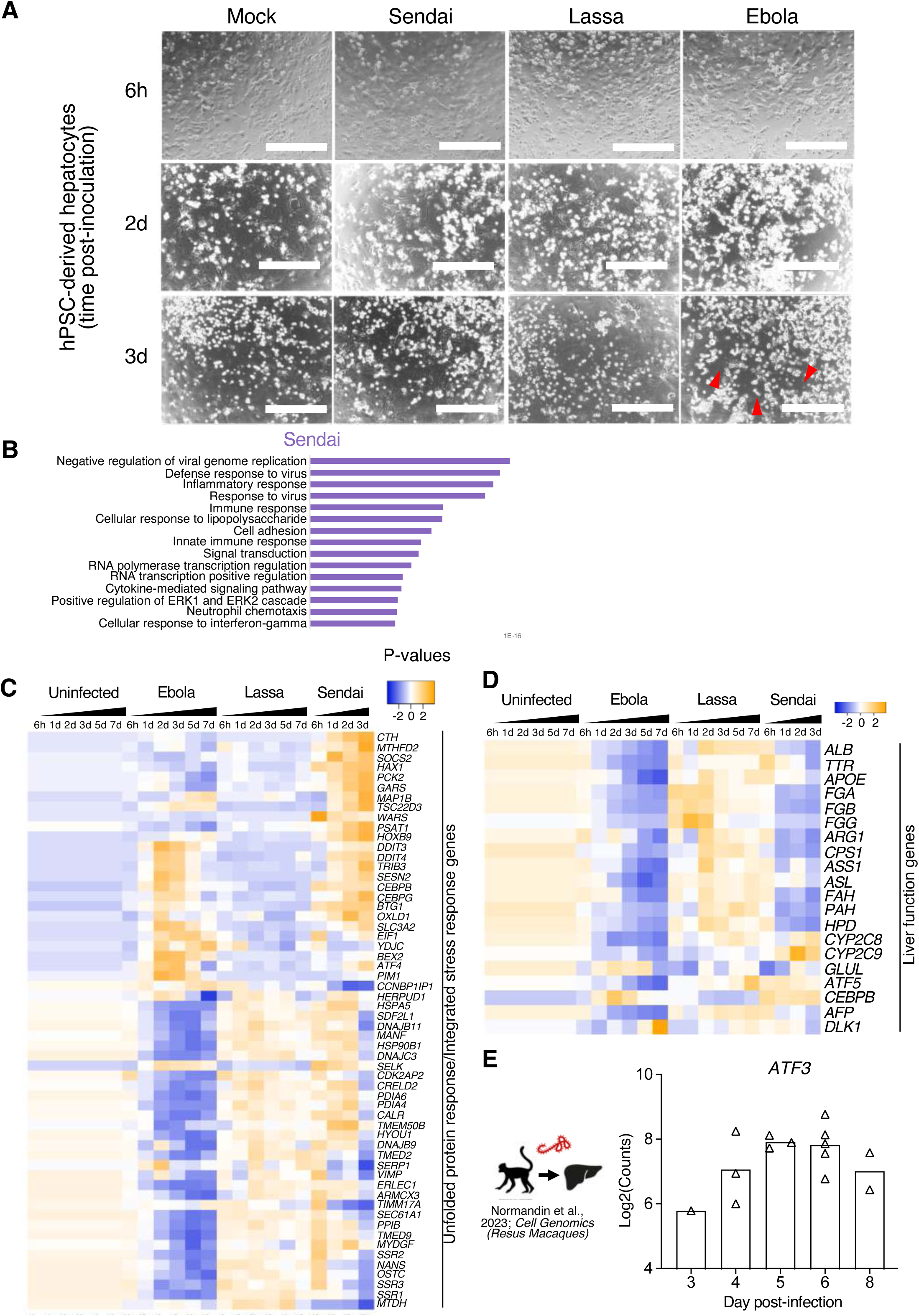
Transcriptional effects of Ebola, Lassa, and Sendai virus on purified hPSC-derived hepatocytes, related to Fig. 7. A) Brightfield images of uninfected hPSC-derived hepatocytes, or those infected by Ebola, Lassa, or Sendai virus. Red arrows indicate overt cell death after Ebola virus infection. Scale bar, 400μm. B) RNA-seq comparison of Sendai virus-vs. mock-infected hPSC-derived hepatocytes to determine gene ontology (GO) terms associated with Sendai virus-upregulated genes. C) Bulk population RNA-seq of uninfected hPSC-derived hepatocytes, or those infected by Ebola, Lassa, and Sendai virus, showing expression of genes linked to the unfolded protein response and integrated stress response^168, 169^. D) Bulk population RNA-seq of uninfected hPSC-derived hepatocytes, or those infected by Ebola, Lassa, and Sendai virus, showing expression of liver function genes.

## INVENTORY OF SUPPLEMENTARY INFORMATION

**Table S1.** Bulk-population RNA-seq of hPSC differentiation into hepatocytes.

**Table S2.** Flow cytometry antibody high-throughput screen of H1 day 18 hPSC-derived hepatocytes that are FAH-Clover-negative and FAH-Clover-positive.

**Table S3.** Differential gene expression analysis of day 18 hPSC-derived hepatocytes vs. H1 hPSC (p < 0.01 and log2FC > 1.5).

**Table S4.** Differential gene expression analysis of day 19 unselected vs. day 19 selected hPSC-derived hepatocytes (p < 0.01).

**Table S5.** Bulk-population RNA-seq of “metabolically-selected” hPSC-derived hepatocytes, HepG2, HuH7, and adult primary human hepatocytes treated and not treated with Poly(I:C). **Table S6.** Bulk-population RNA-seq of cultured adult human hepatocytes and human fetal hepatocytes.

**Table S7.** Critical Reagent or Resource.

**Table S8.** Quantitative PCR primers were used in this study.

**Table S9.** Bulk-population RNA-seq of Ebola, Lassa, and Sendai infected vs. uninfected metabolically-selected hPSC-derived hepatocytes.

## RAW FILES DEPOSITED TO NCBI SRA

- Single-cell RNA-seq of hPSCs and hPSCs differentiating into liver cells at 1, 2, 3, 6, 12, 18 days of differentiation, primary adult hepatocytes, HepG2, and HuH-7 liver cancer cell lines (fastq files)
- Bulk-population RNA-seq of hPSCs and hPSCs differentiating into liver cells (fastq files)
- Omni-ATAC-seq of hPSCs and hPSCs differentiating into liver cells (fastq files)
- Bulk-population RNA-seq of Poly(I:C)-treated vs. mock treatment of hPSC-derived hepatocytes (fastq files)
- Bulk-population RNA-seq fastq files of Ebola, Lassa, and Sendai virus infected hPSC-derived hepatocytes (fastq files)

## Notes

### Competing Interest Statement

The authors have declared no competing interest.

## REFERENCES

1. CDC & NIH Biosafety in Microbiological and Biomedical Laboratories—6th Edition. (U.S. Department of Health and Human Services, Atlanta, GA; 2020).

2. Kuthyar, S., Anthony, C.L., Fashina, T., Yeh, S. & Shantha, J.G. World Health Organization High Priority Pathogens: Ophthalmic Disease Findings and Vision Health Perspectives. Pathogens 10, 442 (2021).

3. Røttingen, J.-A. et al. New Vaccines against Epidemic Infectious Diseases. The New England Journal of Medicine 376, 610–613 (2017).

4. Kuhn, J. Filoviruses : A Compendium of 40 Years of Epidemiological, Clinical, and Laboratory Studies, Edn. 1st 2008. (Springer Vienna : Imprint: Springer, Vienna; 2008).

5. Feldmann, H., Sprecher, A. & Geisbert, T.W. Ebola. N Engl J Med 382, 1832–1842 (2020).

6. Feldmann, H. & Geisbert, T.W. Ebola haemorrhagic fever. Lancet 377, 849–862 (2011).

7. Jacob, S.T. et al. Ebola virus disease. Nat Rev Dis Primers 6, 13 (2020).

8. Ohimain, E.I. & Silas-Olu, D. The 2013-2016 Ebola virus disease outbreak in West Africa. Curr Opin Pharmacol 60, 360–365 (2021).

9. Gatherer, D. The 2014 Ebola virus disease outbreak in West Africa. J. Gen. Virol. 95, 1619–1624 (2014).

10. Dixon, M.G., Schafer, I.J., Centers for Disease, C. & Prevention Ebola viral disease outbreak--West Africa, 2014. MMWR Morb Mortal Wkly Rep 63, 548-551 (2014).

11. Kiggundu, T. et al. Notes from the Field: Outbreak of Ebola Virus Disease Caused by Sudan ebolavirus - Uganda, August-October 2022. MMWR Morb Mortal Wkly Rep 71, 1457-1459 (2022).

12. Musoke, P. & Bongomin, F. Sudan virus disease outbreak in Uganda in 2022: the case of patient zero. Int J Infect Dis 128, 318–320 (2023).

13. Sibomana, O. & Kubwimana, E. First-ever Marburg virus disease outbreak in Equatorial Guinea and Tanzania: An imminent crisis in West and East Africa. Immun Inflamm Dis 11, e980 (2023).

14. Prescott, J.B. et al. Immunobiology of Ebola and Lassa virus infections. Nature Reviews Immunology 17, 195–207 (2017).

15. Gunther, S. & Lenz, O. Lassa virus. Crit Rev Clin Lab Sci 41, 339–390 (2004).

16. Andersen, K.G. et al. Clinical Sequencing Uncovers Origins and Evolution of Lassa Virus. Cell 162, 738–750 (2015).

17. Caballero, I.S. et al. Lassa and Marburg viruses elicit distinct host transcriptional responses early after infection. BMC Genomics 15, 960 (2014).

18. Paassen, J.v., et al. Acute liver failure, multiorgan failure, cerebral oedema, and activation of proangiogenic and antiangiogenic factors in a case of Marburg haemorrhagic fever. The Lancet Infectious Diseases 12, 635–642 (2012).

19. Geisbert, T.W. et al. Marburg Virus Angola Infection of Rhesus Macaques: Pathogenesis and Treatment with Recombinant Nematode Anticoagulant Protein c2. The Journal of Infectious Diseases 196, S372–S381 (2007).

20. McCormick, J.B. et al. Lassa Virus Hepatitis: a Study of Fatal Lassa Fever in Humans. Am J Tropical Medicine Hyg 35, 401–407 (1986).

21. Martines, R.B., Ng, D.L., Greer, P.W., Rollin, P.E. & Zaki, S.R. Tissue and cellular tropism, pathology and pathogenesis of Ebola and Marburg viruses. The Journal of Pathology 235, 153–174 (2014).

22. Rollin, P.E., Bausch, D.G. & Sanchez, A. Blood Chemistry Measurements and d-Dimer Levels Associated with Fatal and Nonfatal Outcomes in Humans Infected with Sudan Ebola Virus. The Journal of Infectious Diseases 196, S364–S371 (2007).

23. Johnson, K.M., Lange, J.V., Webb, P.A. & Murphy, F.A. Isolation and Partial Characterisation of a New Virus causing Acute Haemorrhagic Fever in Zaire. The Lancet 309, 569–571 (1977).

24. Hensley, L.E. et al. Pathogenesis of lassa fever in cynomolgus macaques. Virol J 8, 205 (2011).

25. Shieh, W.-J. et al. Pathology and Pathogenesis of Lassa Fever: Novel Immunohistochemical Findings in Fatal Cases and Clinico-pathologic Correlation. Clin. Infect. Dis. 74, ciab719-(2021).

26. Walker, D.H. et al. Pathologic and virologic study of fatal Lassa fever in man. Am J Pathol 107, 349–356 (1982).

27. Bechtelsheimer, H., Korb, G. & Gedigk, P. The morphology and pathogenesis of “Marburg virus” hepatitis. Hum Pathol 3, 255–264 (1972).

28. Ellis, D.S., Bowen, E.T., Simpson, D.I. & Stamford, S. Ebola virus: a comparison, at ultrastructural level, of the behaviour of the Sudan and Zaire strains in monkeys. Br J Exp Pathol 59, 584–593 (1978).

29. Ellis, D.S. et al. Ultrastructure of Ebola virus particles in human liver. J Clin Pathol 31, 201–208 (1978).

30. Greenberg, A. et al. Quantification of Viral and Host Biomarkers in the Liver of Rhesus Macaques A Longitudinal Study of Zaire Ebolavirus Strain Kikwit (EBOV/Kik). The American Journal of Pathology 190, 1449–1460 (2020).

31. Vernet, M.A. et al. Clinical, virological, and biological parameters associated with outcomes of Ebola virus infection in Macenta, Guinea. JCI Insight 2, e88864 (2017).

32. Cross, R.W. et al. Natural history of nonhuman primates after conjunctival exposure to Ebola virus. Sci Rep 13, 4175 (2023).

33. Geisbert, T.W. & Hensley, L.E. Ebola virus: new insights into disease aetiopathology and possible therapeutic interventions. Expert Rev Mol Med 6, 1–24 (2004).

34. Jankeel, A. et al. Early Transcriptional Changes within Liver, Adrenal Gland, and Lymphoid Tissues Significantly Contribute to Ebola Virus Pathogenesis in Cynomolgus Macaques. J. Virol. 94 (2020).

35. Woolsey, C. et al. Natural history of Sudan ebolavirus infection in rhesus and cynomolgus macaques. Emerg Microbes Infect 11, 1635–1646 (2022).

36. Carrion, R., Jr., et al. Lassa virus infection in experimentally infected marmosets: liver pathology and immunophenotypic alterations in target tissues. J. Virol. 81, 6482–6490 (2007).

37. Shurtleff, A.C. et al. The impact of regulations, safety considerations and physical limitations on research progress at maximum biocontainment. Viruses 4, 3932–3951 (2012).

38. Hartman, A.L., Dover, J.E., Towner, J.S. & Nichol, S.T. Reverse Genetic Generation of Recombinant Zaire Ebola Viruses Containing Disrupted IRF-3 Inhibitory Domains Results in Attenuated Virus Growth In Vitro and Higher Levels of IRF-3 Activation without Inhibiting Viral Transcription or Replication. J. Virol. 80, 6430–6440 (2006).

39. Hartman, A.L., Ling, L., Nichol, S.T. & Hibberd, M.L. Whole-Genome Expression Profiling Reveals That Inhibition of Host Innate Immune Response Pathways by Ebola Virus Can Be Reversed by a Single Amino Acid Change in the VP35 Protein. J. Virol. 82, 5348–5358 (2008).

40. Logue, J. et al. Ebola Virus Isolation Using Huh-7 Cells has Methodological Advantages and Similar Sensitivity to Isolation Using Other Cell Types and Suckling BALB/c Laboratory Mice. Viruses 11, 161 (2019).

41. Kuzmin, I.V. et al. Innate Immune Responses of Bat and Human Cells to Filoviruses: Commonalities and Distinctions. J Virol 91 (2017).

42. Flint, M. et al. A genome-wide CRISPR screen identifies N-acetylglucosamine-1-phosphate transferase as a potential antiviral target for Ebola virus. Nat Commun 10, 285 (2019).

43. Welch, S.R. et al. Lassa and Ebola virus inhibitors identified using minigenome and recombinant virus reporter systems. Antiviral Res 136, 9–18 (2016).

44. Zhou, B. et al. Haplotype-resolved and integrated genome analysis of the cancer cell line HepG2. Nucleic Acids Research 47, gkz169-(2019).

45. Kasai, F., Hirayama, N., Ozawa, M., Satoh, M. & Kohara, A. HuH-7 reference genome profile: complex karyotype composed of massive loss of heterozygosity. Hum Cell 31, 261–267 (2018).

46. Marozin, S. et al. Inhibition of the IFN-β Response in Hepatocellular Carcinoma by Alternative Spliced Isoform of IFN Regulatory Factor-3. Molecular Therapy 16, 1789–1797 (2008).

47. Li, K., Chen, Z., Kato, N., Gale, M. & Lemon, S.M. Distinct Poly(I-C) and Virus-activated Signaling Pathways Leading to Interferon-β Production in Hepatocytes*. Journal of Biological Chemistry 280, 16739–16747 (2005).

48. Scoon, W.A., et al. Ebola virus infection induces a delayed type I IFN response in bystander cells and the shutdown of key liver genes in human iPSC-derived hepatocytes. Stem Cell Reports (2022).

49. Cotovio, J.P. & Fernandes, T.G. Production of Human Pluripotent Stem Cell-Derived Hepatic Cell Lineages and Liver Organoids: Current Status and Potential Applications. Bioeng 7, 36 (2020).

50. Maepa, S.W. & Ndlovu, H. Advances in generating liver cells from pluripotent stem cells as a tool for modeling liver diseases. Stem Cells 38, 606–612 (2020).

51. Tracy, T.S. et al. Interindividual Variability in Cytochrome P450–Mediated Drug Metabolism. Drug Metabolism and Disposition 44, 343–351 (2016).

52. Telles-Silva, K.A. et al. Applied Hepatic Bioengineering: Modeling the Human Liver Using Organoid and Liver-on-a-Chip Technologies. Frontiers Bioeng Biotechnology 10, 845360 (2022).

53. Ardisasmita, A.I. et al. A comprehensive transcriptomic comparison of hepatocyte model systems improves selection of models for experimental use. Commun Biol 5, 1094 (2022).

54. Fowler, J.L., Ang, L.T. & Loh, K.M. A critical look: Challenges in differentiating human pluripotent stem cells into desired cell types and organoids. Wiley Interdisciplinary Reviews: Developmental Biology 113, 891-823 (2019).

55. Thomson, J.A. et al. Embryonic Stem Cell Lines Derived from Human Blastocysts. Science 282, 1145–1147 (1998).

56. Takahashi, K. et al. Induction of Pluripotent Stem Cells from Adult Human Fibroblasts by Defined Factors. Cell 131, 861–872 (2007).

57. Tan, A.K.Y., Loh, K.M. & Ang, L.T. Evaluating the regenerative potential and functionality of human liver cells in mice. Differentiation 98, 25–34 (2017).

58. Levenstein, M.E. et al. Basic fibroblast growth factor support of human embryonic stem cell self-renewal. Stem Cells 24, 568–574 (2006).

59. Takebe, T. et al. Vascularized and functional human liver from an iPSC-derived organ bud transplant. Nature 499, 481–484 (2013).

60. Ogawa, S. et al. Three-dimensional culture and cAMP signaling promote the maturation of human pluripotent stem cell-derived hepatocytes. Development 140, 3285–3296 (2013).

61. Carpentier, A. et al. Engrafted human stem cell–derived hepatocytes establish an infectious HCV murine model. Journal of Clinical Investigation 124, 4953–4964 (2014).

62. Si Tayeb, K., et al. Highly efficient generation of human hepatocyte–like cells from induced pluripotent stem cells. Hepatology 51, 297–305 (2010).

63. Zhao, D. et al. Promotion of the efficient metabolic maturation of human pluripotent stem cell-derived hepatocytes by correcting specification defects. Cell Research 23, 157–161 (2012).

64. Touboul, T. et al. Generation of functional hepatocytes from human embryonic stem cells under chemically defined conditions that recapitulate liver development. Hepatology 51, 1754–1765 (2010).

65. Sumi, T., Tsuneyoshi, N., Nakatsuji, N. & Suemori, H. Defining early lineage specification of human embryonic stem cells by the orchestrated balance of canonical Wnt/ -catenin, Activin/Nodal and BMP signaling. Development 135, 2969–2979 (2008).

66. Yanagida, A., Ito, K., Chikada, H., Nakauchi, H. & Kamiya, A. An In Vitro Expansion System for Generation of Human iPS Cell-Derived Hepatic Progenitor-Like Cells Exhibiting a Bipotent Differentiation Potential. PLoS ONE 8, e67541 (2013).

67. Gadue, P., Huber, T.L., Paddison, P.J. & Keller, G.M. Wnt and TGF-β signaling are required for the induction of an in vitro model of primitive streak formation using embryonic stem cells. Proceedings of the National Academy of Sciences 103, 16806–16811 (2006).

68. Gouon-Evans, V. et al. BMP-4 is required for hepatic specification of mouse embryonic stem cell–derived definitive endoderm. Nature Biotechnology 24, 1402–1411 (2006).

69. Ang, L.T. et al. A Roadmap for Human Liver Differentiation from Pluripotent Stem Cells. Cell Reports 22, 2190–2205 (2018).

70. Cai, J. et al. Directed differentiation of human embryonic stem cells into functional hepatic cells. Hepatology 45, 1229–1239 (2007).

71. Agarwal, S., Holton, K.L. & Lanza, R. Efficient Differentiation of Functional Hepatocytes from Human Embryonic Stem Cells. Stem Cells 26, 1117–1127 (2008).

72. Haridass, D. et al. Repopulation Efficiencies of Adult Hepatocytes, Fetal Liver Progenitor Cells, and Embryonic Stem Cell-Derived Hepatic Cells in Albumin-Promoter-Enhancer Urokinase-Type Plasminogen Activator Mice. The American Journal of Pathology 175, 1483–1492 (2009).

73. Han, S. Generation of Functional Hepatic Cells from Pluripotent Stem Cells. Journal of Stem Cell Research and Therapy 1, 1–7 (2012).

74. Song, Z. et al. Efficient generation of hepatocyte-like cells from human induced pluripotent stem cells. Cell Research 19, 1233–1242 (2009).

75. Shiraki, N. et al. Differentiation of mouse and human embryonic stem cells into hepatic lineages. Genes to Cells 13, 731–746 (2008).

76. Avior, Y. et al. Microbial-derived lithocholic acid and vitamin K2 drive the metabolic maturation of pluripotent stem cells-derived and fetal hepatocytes. Hepatology 62, 265–278 (2015).

77. Carpentier, A. et al. Hepatic differentiation of human pluripotent stem cells in miniaturized format suitable for high-throughput screen. Stem Cell Research 16, 640–650 (2016).

78. Tolosa, L. et al. Transplantation of hESC-derived hepatocytes protects mice from liver injury. Stem Cell Research & Therapy 6, 246 (2015).

79. Nagamoto, Y. et al. Transplantation of a human iPSC-derived hepatocyte sheet increases survival in mice with acute liver failure. Journal of Hepatology 64, 1068–1075 (2016).

80. Ma, H. et al. The nuclear receptor THRB facilitates differentiation of human PSCs into more mature hepatocytes. Cell Stem Cell 29, 795–809 e711 (2022).

81. Loh, K.M. et al. Efficient Endoderm Induction from Human Pluripotent Stem Cells by Logically Directing Signals Controlling Lineage Bifurcations. Cell Stem Cell 14, 237–252 (2014).

82. Tohyama, S., Tanosaki, S., Someya, S., Fujita, J. & Fukuda, K. Manipulation of Pluripotent Stem Cell Metabolism for Clinical Application. Curr Stem Cell Reports 3, 28–34 (2017).

83. Tohyama, S. et al. Distinct Metabolic Flow Enables Large-Scale Purification of Mouse and Human Pluripotent Stem Cell-Derived Cardiomyocytes. Cell Stem Cell 12, 127–137 (2013).

84. Lin, B. et al. Culture in Glucose-Depleted Medium Supplemented with Fatty Acid and 3,3′,5-Triiodo-l-Thyronine Facilitates Purification and Maturation of Human Pluripotent Stem Cell-Derived Cardiomyocytes. Frontiers in endocrinology 8, E1848 (2017).

85. Alexander, P.B., Wang, J. & McKnight, S.L. Targeted killing of a mammalian cell based upon its specialized metabolic state. Proc Natl Acad Sci U S A 108, 15828–15833 (2011).

86. Taya, Y. et al. Depleting dietary valine permits nonmyeloablative mouse hematopoietic stem cell transplantation. Science 354, 1152–1155 (2016).

87. Wang, J. et al. Dependence of mouse embryonic stem cells on threonine catabolism. Science 325, 435–439 (2009).

88. Han, H.-S., Kang, G., Kim, J.S., Choi, B.H. & Koo, S.-H. Regulation of glucose metabolism from a liver-centric perspective. Exp Mol Medicine 48, e218–e218 (2016).

89. Rui, L. Energy Metabolism in the Liver. Compr Physiol 4, 177–197 (2017).

90. Adeva-Andany, M.M., Pérez-Felpete, N., Fernández-Fernández, C., Donapetry-García, C. & Pazos-García, C. Liver glucose metabolism in humans. Bioscience Rep 36, e00416 (2016).

91. Klover, P.J. & Mooney, R.A. Hepatocytes: critical for glucose homeostasis. Int J Biochem Cell Biology 36, 753–758 (2004).

92. Peters, D.T. et al. Asialoglycoprotein receptor 1 is a specific cell-surface marker for isolating hepatocytes derived from human pluripotent stem cells. Development 143, 1475–1481 (2016).

93. Zheng, G.X.Y. et al. Massively parallel digital transcriptional profiling of single cells. Nature Communications 8, 14049 (2017).

94. Corces, M.R. et al. An improved ATAC-seq protocol reduces background and enables interrogation of frozen tissues. Nature Methods 14, 959–962 (2017).

95. Kanai-Azuma, M. et al. Depletion of definitive gut endoderm in Sox17-null mutant mice. Development 129, 2367 - 2379 (2002).

96. Engert, S., Liao, W.P., Burtscher, I. & Lickert, H. Sox17-2A-iCre: A knock-in mouse line expressing Cre recombinase in endoderm and vascular endothelial cells. Genesis 47, 603–610 (2009).

97. Engert, S., Burtscher, I., Kalali, B., Gerhard, M. & Lickert, H. The Sox17CreERT2 knock-in mouse line displays spatiotemporal activation of Cre recombinase in distinct Sox17 lineage progenitors. Genesis 51, 793–802 (2013).

98. Tam, P.P.L. & Loebel, D.A.F. Gene function in mouse embryogenesis: get set for gastrulation. Nature Reviews Genetics 8, 368–381 (2007).

99. Lu, C.C., Robertson, E.J. & Brennan, J. The mouse frizzled 8 receptor is expressed in anterior organizer tissues. Gene Expr Patterns 4, 569–572 (2004).

100. Blum, M. et al. Gastrulation in the mouse: the role of the homeobox gene goosecoid. Cell 69, 1097–1106 (1992).

101. Shawlot, W., Deng, J.M. & Behringer, R.R. Expression of the mouse cerberus-related gene, Cerr1, suggests a role in anterior neural induction and somitogenesis. Proc Natl Acad Sci USA 95, 6198-6203 (1998).

102. Biben, C. et al. Murine cerberus homologue mCer-1: a candidate anterior patterning molecule. Developmental Biology 194, 135–151 (1998).

103. Lüdtke, T.H.-W., Christoffels, V.M., Petry, M. & Kispert, A. Tbx3 promotes liver bud expansion during mouse development by suppression of cholangiocyte differentiation. Hepatology 49, 969–978 (2009).

104. Guz, Y. et al. Expression of murine STF-1, a putative insulin gene transcription factor, in beta cells of pancreas, duodenal epithelium and pancreatic exocrine and endocrine progenitors during ontogeny. Development 121, 11–18 (1995).

105. Chawengsaksophak, K., Graaff, W.d., Rossant, J., Deschamps, J. & Beck, F. Cdx2 is essential for axial elongation in mouse development. Proceedings of the National Academy of Sciences 101, 7641 - 7645 (2004).

106. Beck, F., Erler, T., Russell, A. & James, R. Expression of Cdx-2 in the mouse embryo and placenta: possible role in patterning of the extra-embryonic membranes. Dev Dyn 204, 219–227 (1995).

107. Zorn, A.M. & Wells, J.M. Vertebrate endoderm development and organ formation. Annu Rev Cell Dev Biol 25, 221–251 (2009).

108. Aihara, E., Engevik, K.A. & Montrose, M.H. Trefoil Factor Peptides and Gastrointestinal Function. Annual Review of Physiology 79, 357–380 (2015).

109. Wang, Y. et al. Single-cell transcriptome analysis reveals differential nutrient absorption functions in human intestine. J. Exp. Med. 217, e20191130 (2019).

110. Asada, R. et al. The Endoplasmic Reticulum Stress Transducer OASIS Is involved in the Terminal Differentiation of Goblet Cells in the Large Intestine*. Journal of Biological Chemistry 287, 8144–8153 (2012).

111. Egozi, A. et al. Insulin is expressed by enteroendocrine cells during human fetal development. Nature Medicine 27, 2104–2107 (2021).

112. Engelstoft, M.S., Egerod, K.L., Lund, M.L. & Schwartz, T.W. Enteroendocrine cell types revisited. Curr. Opin. Pharm. 13, 912–921 (2013).

113. Stanger, B.Z., Datar, R., Murtaugh, L.C. & Melton, D.A. Direct regulation of intestinal fate by Notch. Proceedings of the National Academy of Sciences 102, 12443–12448 (2005).

114. Jensen, J. et al. Control of endodermal endocrine development by Hes-1. Nature Genetics 24, 36–44 (2000).

115. Es, J.H.v., et al. Notch/γ-secretase inhibition turns proliferative cells in intestinal crypts and adenomas into goblet cells. Nature 435, 959–963 (2005).

116. Kvittingen, E.A. Hereditary tyrosinemia type I--an overview. Scand J Clin Lab Invest Suppl 184, 27–34 (1986).

117. Fagerberg, L. et al. Analysis of the Human Tissue-specific Expression by Genome-wide Integration of Transcriptomics and Antibody-based Proteomics. Molecular & Cellular Proteomics 13, 397–406 (2014).

118. Aizarani, N. et al. A human liver cell atlas reveals heterogeneity and epithelial progenitors. Nature 572, 1–6 (2019).

119. Tohyama, S. et al. Glutamine Oxidation Is Indispensable for Survival of Human Pluripotent Stem Cells. Cell Metabolism 23, 663–674 (2016).

120. Tomizawa, M. et al. Survival of Primary Human Hepatocytes and Death of Induced Pluripotent Stem Cells in Media Lacking Glucose and Arginine. PLoS ONE 8, e71897 (2013).

121. Zhou, T. et al. The role of PEG3 in the occurrence and prognosis of colon cancer. Onco Targets Ther 12, 6001–6012 (2019).

122. Xie, T. et al. PEG10 as an oncogene: expression regulatory mechanisms and role in tumor progression. Cancer Cell Int 18, 112 (2018).

123. Tabula Muris, C., et al. Single-cell transcriptomics of 20 mouse organs creates a Tabula Muris. Nature 562, 367–372 (2018).

124. Naraharisetti, S.B. et al. Human Liver Expression of CYP2C8: Gender, Age, and Genotype Effects. Drug Metabolism and Disposition 38, 889–893 (2010).

125. Michaels, S. & Wang, M.Z. The Revised Human Liver Cytochrome P450 “Pie”: Absolute Protein Quantification of CYP4F and CYP3A Enzymes Using Targeted Quantitative Proteomics. Drug Metabolism and Disposition 42, 1241–1251 (2014).

126. Hart, S.N., Cui, Y., Klaassen, C.D. & Zhong, X.b. Three Patterns of Cytochrome P450 Gene Expression during Liver Maturation in Mice. Drug Metabolism and Disposition 37, 116–121 (2008).

127. Blake, M.J., Castro, L., Leeder, J.S. & Kearns, G.L. Ontogeny of drug metabolizing enzymes in the neonate. Seminars in Fetal and Neonatal Medicine 10, 123–138 (2005).

128. Hines, R.N. Ontogeny of human hepatic cytochromes P450. J. Biochem. Mol. Toxicol. 21, 169–175 (2007).

129. Nitsch, D., Boshart, M. & Schutz, G. Activation of the tyrosine aminotransferase gene is dependent on synergy between liver-specific and hormone-responsive elements. Proc Natl Acad Sci U S A 90, 5479–5483 (1993).

130. West, G. et al. Key differences between apoC-III regulation and expression in intestine and liver. Biochem. Biophys. Res. Commun. 491, 747–753 (2017).

131. Haddad, I.A., Ordovas, J.M., Fitzpatrick, T. & Karathanasis, S.K. Linkage, evolution, and expression of the rat apolipoprotein A-I, C-III, and A-IV genes. J. Biol. Chem. 261, 13268–13277 (1986).

132. Jungermann, K. Zonation of metabolism and gene expression in liver. Histochemistry and Cell Biology 103, 81–91 (1995).

133. Gebhardt, R. Metabolic zonation of the liver: regulation and implications for liver function. Pharmacology and Therapeutics 53, 275–354 (1992).

134. Colnot, S. & Perret, C. Liver Zonation, Vol. 5 7-16 (Springer US, Boston, MA; 2011).

135. Pek, N.M.Q., Liu, K.J., Nichane, M. & Ang, L.T. Controversies Surrounding the Origin of Hepatocytes in Adult Livers and the in Vitro Generation or Propagation of Hepatocytes. Cellular and Molecular Gastroenterology and Hepatology, 1–18 (2020).

136. Alves-Bezerra, M. & Cohen, D.E. Triglyceride Metabolism in the Liver. Compr Physiol 8, 1–8 (2017).

137. Azuma, H. et al. Robust expansion of human hepatocytes in Fah−/−/Rag2−/−/Il2rg−/− mice. Nature Biotechnology 25, 903–910 (2007).

138. Smith, D.D. & Campbell, J.W. Distribution of glutamine synthetase and carbamoyl-phosphate synthetase I in vertebrate liver. Proceedings of the National Academy of Sciences 85, 160–164 (1988).

139. Ben-Moshe, S. et al. Spatial sorting enables comprehensive characterization of liver zonation. Nature Metabolism, 1–17 (2019).

140. Halpern, K.B. et al. Single-cell spatial reconstruction reveals global division of labour in the mammalian liver. Nature 542, 352–356 (2017).

141. Lazear, H.M., Schoggins, J.W. & Diamond, M.S. Shared and Distinct Functions of Type I and Type III Interferons. Immunity 50, 907–923 (2019).

142. Alexopoulou, L., Holt, A.C., Medzhitov, R. & Flavell, R.A. Recognition of double-stranded RNA and activation of NF-κB by Toll-like receptor 3. Nature 413, 732–738 (2001).

143. Honda, K. et al. Selective contribution of IFN-α/β signaling to the maturation of dendritic cells induced by double-stranded RNA or viral infection. Proceedings of the National Academy of Sciences 100, 10872–10877 (2003).

144. Shaw, A.E. et al. Fundamental properties of the mammalian innate immune system revealed by multispecies comparison of type I interferon responses. PLoS Biol. 15, e2004086 (2017).

145. Yi, J. et al. Large tumor suppressor homologs 1 and 2 regulate mouse liver progenitor cell proliferation and maturation through antagonism of the coactivators YAP and TAZ. Hepatology 64, 1757–1772 (2016).

146. Yamamoto, J., Udono, M., Miura, S., Sekiya, S. & Suzuki, A. Cell Aggregation Culture Induces Functional Differentiation of Induced Hepatocyte-like Cells through Activation of Hippo Signaling. Cell Rep 25, 183–198 (2018).

147. Driskill, J.H. & Pan, D. The Hippo Pathway in Liver Homeostasis and Pathophysiology. Annu Rev Pathol 16, 299–322 (2021).

148. Sun, P. et al. Maintenance of Primary Hepatocyte Functions In Vitro by Inhibiting Mechanical Tension-Induced YAP Activation. Cell Rep 29, 3212–3222 e3214 (2019).

149. Hoenen, T., Groseth, A., Callison, J., Takada, A. & Feldmann, H. A novel Ebola virus expressing luciferase allows for rapid and quantitative testing of antivirals. Antiviral Res 99, 207–213 (2013).

150. Kainulainen, M.H. et al. Recombinant Sudan virus and evaluation of humoral cross-reactivity between Ebola and Sudan virus glycoproteins after infection or rVSV-DeltaG-ZEBOV-GP vaccination. Emerg Microbes Infect 12, 2265660 (2023).

151. Albarino, C.G., Wiggleton Guerrero, L., Chakrabarti, A.K. & Nichol, S.T. Transcriptional analysis of viral mRNAs reveals common transcription patterns in cells infected by five different filoviruses. PLoS One 13, e0201827 (2018).

152. Rosenke, K. et al. Use of Favipiravir to Treat Lassa Virus Infection in Macaques. Emerg Infect Dis 24, 1696–1699 (2018).

153. Bushmaker, T. et al. Limited benefit of post-exposure prophylaxis with VSV-EBOV in Ebola virus-infected rhesus macaques. J Infect Dis (2023).

154. Butler, S.L. et al. The antigen for Hep Par 1 antibody is the urea cycle enzyme carbamoyl phosphate synthetase 1. Lab Invest 88, 78–88 (2008).

155. López, C.B., Yount, J.S., Hermesh, T. & Moran, T.M. Sendai virus infection induces efficient adaptive immunity independently of type I interferons. Journal of Virology 80, 4538–4545 (2006).

156. Reid, S.P. et al. Ebola virus VP24 binds karyopherin alpha1 and blocks STAT1 nuclear accumulation. J. Virol. 80, 5156–5167 (2006).

157. Basler, C.F. Innate immune evasion by filoviruses. Virology 479-480, 122-130 (2015).

158. Misasi, J. & Sullivan, N.J. Camouflage and misdirection: the full-on assault of ebola virus disease. Cell 159, 477–486 (2014).

159. Messaoudi, I., Amarasinghe, G.K. & Basler, C.F. Filovirus pathogenesis and immune evasion: insights from Ebola virus and Marburg virus. Nat. Rev. Microbiol. 13, 663–676 (2015).

160. Meyer, B. & Ly, H. Inhibition of Innate Immune Responses Is Key to Pathogenesis by Arenaviruses. J. Virol. 90, 3810–3818 (2016).

161. Murphy, H. & Ly, H. Understanding Immune Responses to Lassa Virus Infection and to Its Candidate Vaccines. Vaccines (Basel) 10 (2022).

162. Garry, R.F. Lassa fever - the road ahead. Nat. Rev. Microbiol. 21, 87–96 (2023).

163. Jho, E.H. et al. Wnt/beta-catenin/Tcf signaling induces the transcription of Axin2, a negative regulator of the signaling pathway. Mol. Cell. Biol. 22, 1172–1183 (2002).

164. Huggins, I.J. et al. The WNT target SP5 negatively regulates WNT transcriptional programs in human pluripotent stem cells. Nat Commun 8, 1034 (2017).

165. Yang, J. et al. Beta-catenin signaling in murine liver zonation and regeneration: A Wnt-Wnt situation! Hepatology 60, 964–976 (2014).

166. Pakos-Zebrucka, K. et al. The integrated stress response. EMBO Rep 17, 1374–1395 (2016).

167. Costa-Mattioli, M. & Walter, P. The integrated stress response: From mechanism to disease. Science 368 (2020).

168. Replogle, J.M. et al. Mapping information-rich genotype-phenotype landscapes with genome-scale Perturb-seq. Cell 185, 2559–2575 e2528 (2022).

169. Adamson, B. et al. A Multiplexed Single-Cell CRISPR Screening Platform Enables Systematic Dissection of the Unfolded Protein Response. Cell 167, 1867–1882.e1821 (2016).

170. Normandin, E. et al. Natural history of Ebola virus disease in rhesus monkeys shows viral variant emergence dynamics and tissue-specific host responses. Cell Genom 3, 100440 (2023).

171. Nell, P. et al. Identification of an FXR-modulated liver-intestine hybrid state in iPSC-derived hepatocyte-like cells. J Hepatol 77, 1386–1398 (2022).

172. Belanger, M., Allaman, I. & Magistretti, P.J. Brain energy metabolism: focus on astrocyte-neuron metabolic cooperation. Cell Metab 14, 724–738 (2011).

173. Stumvoll, M. et al. Uptake and release of glucose by the human kidney. Postabsorptive rates and responses to epinephrine. J. Clin. Invest. 96, 2528–2533 (1995).

174. Rosen, E.D. & Spiegelman, B.M. Adipocytes as regulators of energy balance and glucose homeostasis. Nature 444, 847–853 (2006).

175. Drubin, D.G. & Hyman, A.A. Stem cells: the new “model organism”. Molecular Biology of the Cell 28, 1409–1411 (2017).

176. Ang, L.T. et al. Generating human artery and vein cells from pluripotent stem cells highlights the arterial tropism of Nipah and Hendra viruses. Cell 185, 2523–2541 e2530 (2022).

177. Fowler, J.L., ANG, L.T. & Loh, K.M. A critical look: Challenges in differentiating human pluripotent stem cells into desired cell types and organoids. Wiley Interdisciplinary Reviews: Developmental Biology 9, 891 (2020).

178. Sharma, N. & Cappell, M.S. Gastrointestinal and Hepatic Manifestations of Ebola Virus Infection. Dig. Dis. Sci. 60, 2590–2603 (2015).

179. Baseler, L., Chertow, D.S., Johnson, K.M., Feldmann, H. & Morens, D.M. The Pathogenesis of Ebola Virus Disease. Annu Rev Pathol 12, 387–418 (2017).

180. Schulze, R.J., Schott, M.B., Casey, C.A., Tuma, P.L. & McNiven, M.A. The cell biology of the hepatocyte: A membrane trafficking machine. J Cell Biol 218, 2096–2112 (2019).

181. Spengler, J.R. et al. Severity of Disease in Humanized Mice Infected With Ebola Virus or Reston Virus Is Associated With Magnitude of Early Viral Replication in Liver. J. Infect. Dis. 217, 58–63 (2017).

182. Escudero-Perez, B. et al. Comparative pathogenesis of Ebola virus and Reston virus infection in humanized mice. JCI Insight 4 (2019).

183. Sung, P.S. & Shin, E.C. Interferon Response in Hepatitis C Virus-Infected Hepatocytes: Issues to Consider in the Era of Direct-Acting Antivirals. Int J Mol Sci 21 (2020).

184. Ozga, A.J., Chow, M.T. & Luster, A.D. Chemokines and the immune response to cancer. Immunity 54, 859–874 (2021).

185. Peters, C.J., Liu, C.T., Anderson, G.W., Jr., Morrill, J.C. & Jahrling, P.B. Pathogenesis of viral hemorrhagic fevers: Rift Valley fever and Lassa fever contrasted. Rev Infect Dis 11 **Suppl 4**, S743–749 (1989).

186. Cross, R.W. et al. Comparison of the Pathogenesis of the Angola and Ravn Strains of Marburg Virus in the Outbred Guinea Pig Model. J. Infect. Dis. 212 **Suppl 2**, S258–270 (2015).

187. Lavender, K.J. et al. Pathogenicity of Ebola and Marburg Viruses Is Associated With Differential Activation of the Myeloid Compartment in Humanized Triple Knockout-Bone Marrow, Liver, and Thymus Mice. J. Infect. Dis. 218, S409–S417 (2018).

188. Muhlberger, E., Weik, M., Volchkov, V.E., Klenk, H.D. & Becker, S. Comparison of the transcription and replication strategies of marburg virus and Ebola virus by using artificial replication systems. J. Virol. 73, 2333–2342 (1999).

189. Edwards, M.R. et al. Differential Regulation of Interferon Responses by Ebola and Marburg Virus VP35 Proteins. Cell Reports 14, 1632–1640 (2016).

190. Guito, J.C., Albarino, C.G., Chakrabarti, A.K. & Towner, J.S. Novel activities by ebolavirus and marburgvirus interferon antagonists revealed using a standardized in vitro reporter system. Virology 501, 147–165 (2017).

191. Richmond, J.K. & Baglole, D.J. Lassa fever: epidemiology, clinical features, and social consequences. BMJ 327, 1271–1275 (2003).

192. Hawman, D.W. & Feldmann, H. Crimean-Congo haemorrhagic fever virus. Nat. Rev. Microbiol. 21, 463–477 (2023).

193. Lindquist, M.E. et al. Exploring Crimean-Congo Hemorrhagic Fever Virus-Induced Hepatic Injury Using Antibody-Mediated Type I Interferon Blockade in Mice. J. Virol. 92 (2018).

194. Rathore, S.S. et al. Crimean-Congo haemorrhagic fever-induced liver injury: A systematic review and meta-analysis. Int. J. Clin. Pract. 75, e14775 (2021).

195. Rodrigues, R., Paranhos-Baccala, G., Vernet, G. & Peyrefitte, C.N. Crimean-Congo hemorrhagic fever virus-infected hepatocytes induce ER-stress and apoptosis crosstalk. PLoS One 7, e29712 (2012).

196. Maiztegui, J.I. et al. Ultrastructural and immunohistochemical studies in five cases of Argentine hemorrhagic fever. J Infect Dis 132, 35–53 (1975).

197. Davis, R.P. et al. Targeting a GFP reporter gene to the MIXL1 locus of human embryonic stem cells identifies human primitive streak-like cells and enables isolation of primitive hematopoietic precursors. Blood 111, 1876–1884 (2008).

198. Choi, H.M.T. et al. Third-generation in situ hybridization chain reaction: multiplexed, quantitative, sensitive, versatile, robust. Development 145, dev165753-165122 (2018).

199. Stuart, T. et al. Comprehensive Integration of Single-Cell Data. Cell 177, 1888–1902.e1821 (2019).

